# Improving stem cell-derived pancreatic islets using single-cell multiome-inferred regulomes

**DOI:** 10.1101/2022.09.25.509355

**Authors:** Han Zhu, Gaowei Wang, Kim-Vy Nguyen-Ngoc, Dongsu Kim, Michael Miller, Georgina Goss, Jenna Kovsky, Austin R. Harrington, Diane Saunders, Rebecca Melton, Alvin C. Powers, Sebastian Preissl, Francesca M. Spagnoli, Kyle J. Gaulton, Maike Sander

**Affiliations:** Department of Pediatrics, University of California San Diego, La Jolla CA, USA; Pediatric Diabetes Research Center, University of California San Diego, La Jolla CA, USA; Center for Epigenomics, Department of Cellular and Molecular Medicine, University of California San Diego, La Jolla, CA, USA; Centre for Gene Therapy and Regenerative Medicine, King’s College London, UK; Division of Diabetes, Endocrinology and Metabolism, Department of Medicine, Vanderbilt University Medical Center, Nashville, USA; Department of Molecular Physiology & Biophysics, Vanderbilt University School of Medicine, Nashville, USA; VA Tennessee Valley Healthcare System, Nashville, USA; Institute for Genomic Medicine, University of California San Diego, La Jolla CA, USA; Department of Cellular and Molecular Medicine, University of California San Diego, La Jolla CA, USA

**Author notes:** Corresponding author: Maike Sander, 9500 Gilman Drive, #0695, Departments of Pediatrics, University of California San Diego. Authors contributed equally to this work (Han Zhu, Gaowei Wang). Institute of Experimental and Clinical Pharmacology and Toxicology, Faculty of Medicine, University of Freiburg, 79104 Freiburg, Germany (Sebastian Preissl).

**Keywords:** human pluripotent stem cells, islets, β-cell, pancreas, CDX2, development, fetal pancreas, single-cell genomics, gene regulatory network, signals, transcription factors, ATAC-seq, RNA-seq

## Abstract

Pancreatic islet cells derived from human pluripotent stem cells hold great promise for modeling and treating diabetes. Differences between stem cell-derived and primary islets remain, but molecular insights to inform improvements are limited. Here, we acquire single-cell transcriptomes and accessible chromatin profiles during *in vitro* islet differentiation and pancreas from childhood and adult donors for comparison. We delineate major cell types, define their regulomes, and describe spatiotemporal gene regulatory relationships between transcription factors. CDX2 emerged as a regulator of enterochromaffin-like cells, which we show resemble a transient, previously unrecognized, CDX2^+^ pre-β-cell population in fetal pancreas, arguing against a proposed non-pancreatic origin. Furthermore, we observe insufficient activation of signal-dependent transcriptional programs during *in vitro* β-cell maturation and identify sex hormones as drivers of β-cell proliferation in childhood. Altogether, our analysis provides a comprehensive understanding of cell fate acquisition in stem cell-derived islets and a framework for manipulating cell identities and maturity.

## Introduction

The ability to generate pancreatic islet-like clusters from human pluripotent stem cells (hPSCs) holds great promise as a cell replacement therapy and *in vitro* disease model for diabetes. Informed by model organism research, current protocols mimic *in vivo* development by stepwise exposure of hPSCs to growth factors and small molecules (Balboa et al., 2022; Hogrebe et al., 2020; Nair et al., 2019; Nostro et al., 2015; Pagliuca et al., 2014; Rezania et al., 2014; Velazco-Cruz et al., 2019; Veres et al., 2019). Stem cell-derived islets (SC-islets) are comprised of insulin-producing β-cells, glucagon-producing α-cells, and somatostatin-producing δ-cells akin to the cell types found in islets of the pancreas. SC-islets also contain cell types thought to be pancreas-aberrant, such as cells resembling enterochromaffin cells of the intestine (Balboa et al., 2022; Veres et al., 2019). The field currently lacks methodology to control cell type yields or eliminate unwanted populations. Furthermore, despite improvements in differentiation protocols, *in vitro* SC-β-cells are still functionally immature and respond differently to insulin secretory signals than primary β-cells (Balboa et al., 2022). SC-β-cells acquire a more mature state when exposed to an *in vivo* environment by engraftment (Augsornworawat et al., 2020; Balboa et al., 2022; Nair et al., 2019; Pagliuca et al., 2014; Rezania et al., 2014; Vegas et al., 2016; Velazco-Cruz et al., 2019), suggesting competence of SC-β-cells to respond to maturation signals but absence of these signals during *in vitro* differentiation. In mice and humans, β-cell functional maturation occurs postnatally (Aguayo-Mazzucato et al., 2011; Arda et al., 2016; Henquin and Nenquin, 2018; Jermendy et al., 2011; Otonkoski et al., 1988; Rorsman et al., 1989) and is driven by environmental cues (Wortham and Sander, 2021), including nutrient (Jaafar et al., 2019; Jacovetti et al., 2015; Stolovich-Rain et al., 2015) and circadian (Rakshit et al., 2018) signals. However, a comprehensive understanding of the signals mediating β-cell maturation during postnatal life is still lacking.

Single-cell technologies allow for profiling of individual cells, providing an opportunity for detailed molecular characterization of developmental trajectories and states of individual cell types. Transcriptome analysis at single-cell level has defined signatures of cell populations during SC-islet differentiation and has identified genes differentially expressed between SC-β-cells and primary β-cells (Augsornworawat et al., 2020; Balboa et al., 2022; Veres et al., 2019; Weng et al., 2020). However, gene expression alone provides a limited understanding of the regulatory control features of each cell type or state which is determined by transcription factors (TFs) interacting with gene regulatory elements to enable precise control of gene expression through complex gene regulatory networks (GRNs) (Fleck et al., 2021). Such integrated GRN maps are still missing for SC-islet differentiation and primary islets, hampering progress toward controlling cell fates and maturation states during *in vitro* differentiation. By combining single-cell gene expression and chromatin accessibility profiling, it is possible to infer cell type- and cell state-specific GRNs to gain insight into the transcriptional programs driving cell fate acquisition and cell maturation.

Here, we build GRNs from single-cell transcriptome and chromatin accessibility profiling data acquired across an SC-islet developmental time course and from primary childhood and adult donor islets. We infer TFs defining cell identities, connect TFs to downstream target genes, and predict upstream signaling factors. A major finding is that enterochromaffin-like cells (SC-ECs) produced during SC-islet differentiation resemble a pre-β-cell population in the fetal pancreas, suggesting a pancreatic rather than intestinal origin. Deletion of the SC-EC regulator *CDX2* in hPSCs supports a close lineage relationship of SC-ECs and SC-β-cells. By comparing regulatory programs of SC-β-cells and primary β-cells during postnatal maturation, we comprehensively identify candidate signaling pathways involved in β-cell maturation and describe signaling pathways insufficiently activated in SC-β-cells. Together, the established GRNs provide a roadmap for understanding and manipulating islet cell differentiation from stem cells.

## Results

### Chromatin accessibility and gene expression during SC-islet differentiation

The differentiation of pancreatic endocrine cells from hPSCs proceeds through several stages, resulting in the formation of insulin^+^ β-cells, glucagon^+^ α-cells, and somatostatin^+^ δ-cells at the immature SC-islet stage (**Figure 1A** and **Figure S1A**). Glucose-stimulated insulin secretion is acquired during an additional ∼2-week period of SC-islet maturation (**Figure S1B**). To characterize the gene regulatory programs governing SC-islet differentiation and maturation, we conducted single-nucleus ATAC-sequencing (snATAC-seq) and single-cell RNA-sequencing (scRNA-seq) analysis at the pancreatic progenitor (day, D11), endocrine progenitor (D14), immature SC-islet (D21), and maturing SC-islet cell stage (D32/39; **Figure 1A, Table S1A**). After rigorous quality control (see Methods; **Figure S1C,D**), we obtained chromatin accessibility profiles from 65,255 cells and transcriptomes from 25,686 cells across the four stages. Following dimensionality reduction and visualization with UMAP, we annotated cell populations based on promoter chromatin accessibility or RNA expression using canonical genes (**Figure 1B** and **Figure S1E,F**). We identified ten clusters in both snATAC-seq and scRNA-seq data representing distinct cell populations: two pancreatic progenitor cell populations (PP1 and PP2), distinguished by *NKX6-1* expression; *NEUROG3*^high^ early endocrine progenitors (ENP1); α-like endocrine progenitors (ENP-α, *ARX*+); two late endocrine progenitor populations (ENP2 and ENP3), marked by *LMX1A* and *RFX3*, respectively; and differentiated cell types including α-cells (SC-α, *GCG*^+^), β-cells (SC-β, *INS*^+^ and *IAPP*^+^), and δ-cells (SC-δ, *SST*^+^). We also identified a gut enterochromaffin cell-like SC-EC population (*INS*^+^ and *SLC18A1*^+^), previously described in SC-islet cultures (Balboa et al., 2022; Veres et al., 2019). Analysis of cell cluster composition by time point of differentiation showed that cultures at D11 and D14 were mostly comprised of pancreatic and endocrine progenitors, whereas cultures at immature and maturing islet stages predominantly contained differentiated endocrine cell types (**Figure S1G,H**). To integrate chromatin accessibility and gene expression data, we performed canonical correlation analysis (see Methods), which allowed us to match cluster annotations between the two datasets (**Figure 1C**) and to generate in-silico pseudo-cells with matched epigenomic and transcriptomic information. The data integration revealed a larger degree of cell type specificity in gene expression than in chromatin accessibility (**Figure 1D**), suggesting plasticity among the cell populations.

**Figure 1.**
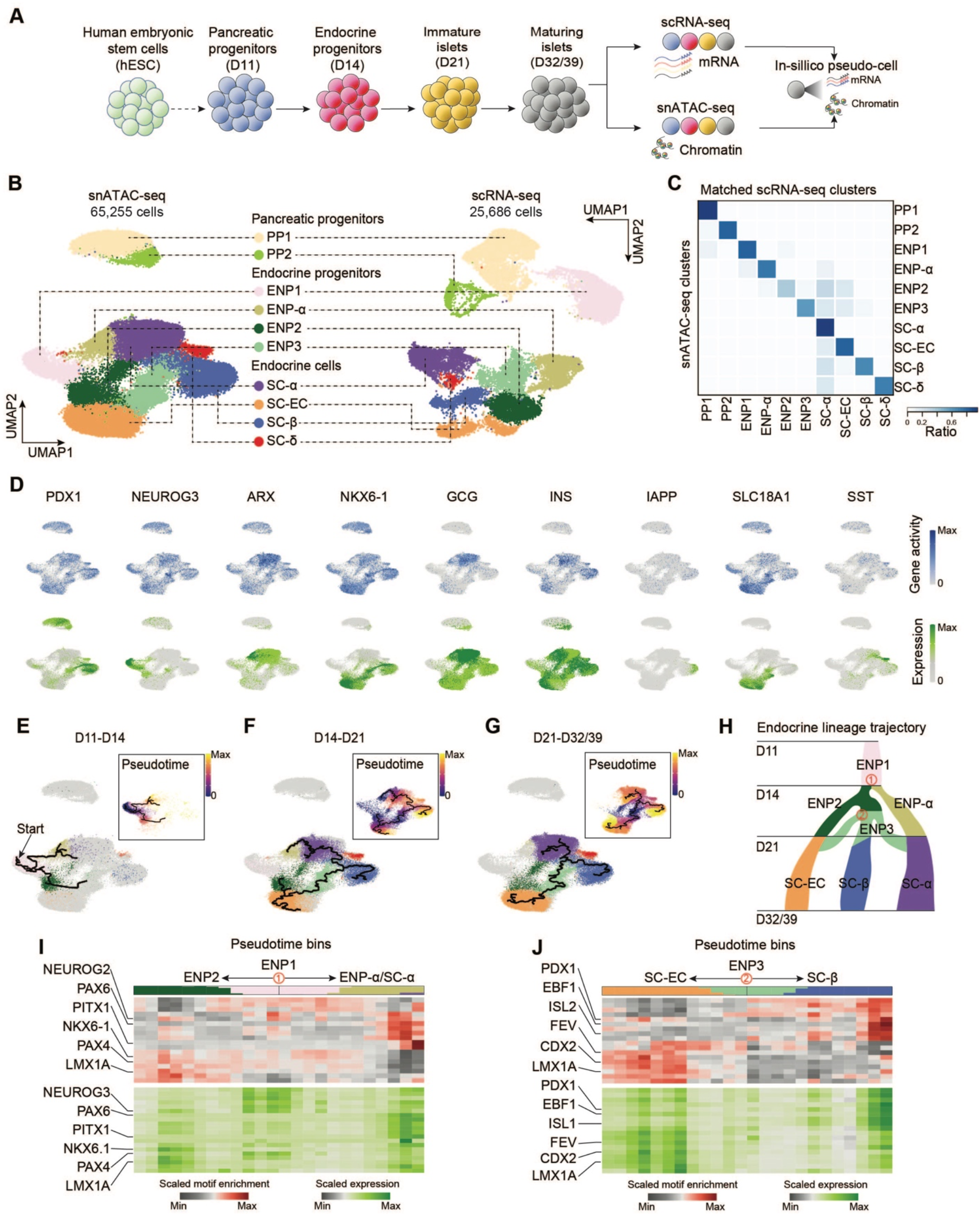
Stem cell islet lineage trajectories based on integrated single-cell chromatin accessibility and transcriptome profiles. (A) Schematic of experimental design. scRNA-seq and snATAC-seq data were obtained during differentiation of islet cells from hESCs at the pancreatic progenitor (day (D) 11), endocrine progenitor (D14), immature islet (D21), and maturing islet (D32 and D39) stage and computationally integrated to generate “pseudo-cells”. (B) UMAP embedding of chromatin accessibility (left) and transcriptome (right) data. Cluster identities were defined by promoter accessibility (snATAC-seq) or expression (scRNA-seq) of marker genes. PP, pancreatic progenitor; ENP, endocrine progenitor; SC-EC, stem cell-derived enterochromaffin cell-like cells. (C) Heatmap showing ratio of cells with identities in scRNA-seq (column) data matching identities in snATAC-seq (row) data. (D) Gene activity (top) and gene expression (bottom) for cell type marker genes. (E-G) Trajectory analysis based on chromatin accessibility, showing trajectories from D11 and D14 (E), D14 and D21 (F), and D21 and D32/39 (G) data with ENP1, ENP2 and ENP3 set as the root, respectively. Cells were color-coded by either cluster identities or pseudotime values (insets). PP1 and PP2 cells were excluded from the analysis. (H) Inferred endocrine lineage trajectory from e-g. Two branch points (in red) were used in analyses in (I) and (J). (I, J) Heatmaps of transcription factor motif enrichment (top) and gene expression (bottom) along pseudotime bins downstream of trajectory branch points in (H). Top bar shows proportion of cell types in each pseudotime bin, using matching colors to cell type annotations in (B).

Given that chromatin accessibility signifies developmental potential beyond cell identity defined by gene expression (Beerman and Rossi, 2015), we inferred lineage relationships between cell populations by trajectory analysis based on chromatin activity using Monocle3 (Cao et al., 2019). To establish the trajectory, we performed analyses of cell populations between two consecutive stages of differentiation (**Figure 1E-G** and **Figure S1I-K**). This analysis identified ENP1 progenitors as a common precursor for all endocrine cell lineages. ENP1 progenitors were predicted to give rise to α-lineage-restricted ENP-α progenitors and ENP2 progenitors that generate SC-ECs as well as ENP3 progenitors with lineage potential for SC-β-cells, SC-ECs, and SC-α-cells (**Figure 1H**). These results indicate that SC-α-cells can be produced from two different progenitor populations, explaining findings from gene expression-based trajectories which have suggested that SC-α-cells can form before or after the specification of the SC-β-cell and SC-EC lineages (Veres et al., 2019; Weng et al., 2020). Overall, this analysis suggests close relatedness of SC-β-cells and SC-ECs.

To identify TFs governing these lineage transitions, we analyzed lineage trajectories for TF binding motif enrichment and expression of TFs corresponding to highly dynamic motifs across the trajectory (see Methods). We focused on two branch points in the lineage tree (**Figure 1H**): (i) separation of the α-cell lineage from the other endocrine cell lineages through generation of ENP-α and ENP2 progenitors from ENP1 progenitors (**Figure 1I** and **Table S2A-C**) and (ii) lineage bifurcation between SC-β-cells and SC-ECs from ENP3 progenitors (**Figure 1J** and **Table S2D-F**). The analysis identified known regulators as well as candidate novel regulators of cell lineage decisions during endocrine cell development. For example, NEUROG3 motif enrichment and expression was highest in ENP1 pan-endocrine progenitors, consistent with its function in endocrine lineage induction (Apelqvist et al., 1999; Gradwohl et al., 2000), whereas PAX6 and PAX4 activity were highest in SC-α- and SC-β-cell precursors (Sander et al., 1997; Sosa-Pineda et al., 1997; St-Onge et al., 1997), respectively (**Figure 1I**). TFs with unknown function in endocrine cell development included PITX1 predicted to specify SC-α-cells and LMX1A with a predicted role in non-α lineage choices (**Figure 1I**). Analysis of the SC-β-cell versus SC-EC lineage branch point confirmed PDX1 as a β-cell regulator (Gao et al., 2014; Holland et al., 2002) and suggested a similar role for EBF1 (**Figure 1J**). Interestingly, the TFs FEV, LMX1A, and CDX2 exhibited motif enrichment and higher expression in SC-ECs compared to ENP3 progenitors or SC-β-cells (**Figure 1J**), indicating roles for FEV, LMX1A, and CDX2 in SC-EC lineage specification.

### Characterization of cell type-specific gene regulatory programs

To comprehensively characterize gene regulatory programs governing SC-islet cell type differentiation, we inferred gene regulatory networks (GRNs) for each cell population, linking TFs to candidate *cis*-regulatory elements (cCREs) and their target genes (see Methods and **Figure 2A**). Briefly, we aggregated cells with the same cell type identity and time point of differentiation for a total of 15 pseudo-bulk bins and linked distal cCREs to target genes based on correlation between cCRE accessibility and gene expression (**Figure S2A-C**). This analysis yielded 65,479 cCRE-gene pairs (FDR<0.15; total of 51,281 cCREs and 11,997 target genes) with each gene being regulated by an average of 5.6 cCREs and each cCRE controlling an average of 1.3 genes (**Figure S2D,E**). Next, to link cCREs to upstream TFs, we performed motif analysis (Schep et al., 2017) and correlated TF expression and cCRE accessibility among TF-cCRE pairs as an indicator of TF-cCRE interaction strength. 266 TFs were predicted to bind the 51,281 cCREs within the GRN (Spearman correlation>0.5) with each TF binding to 1,053 cCREs (**Figure S2F,G**).

**Figure 2.**
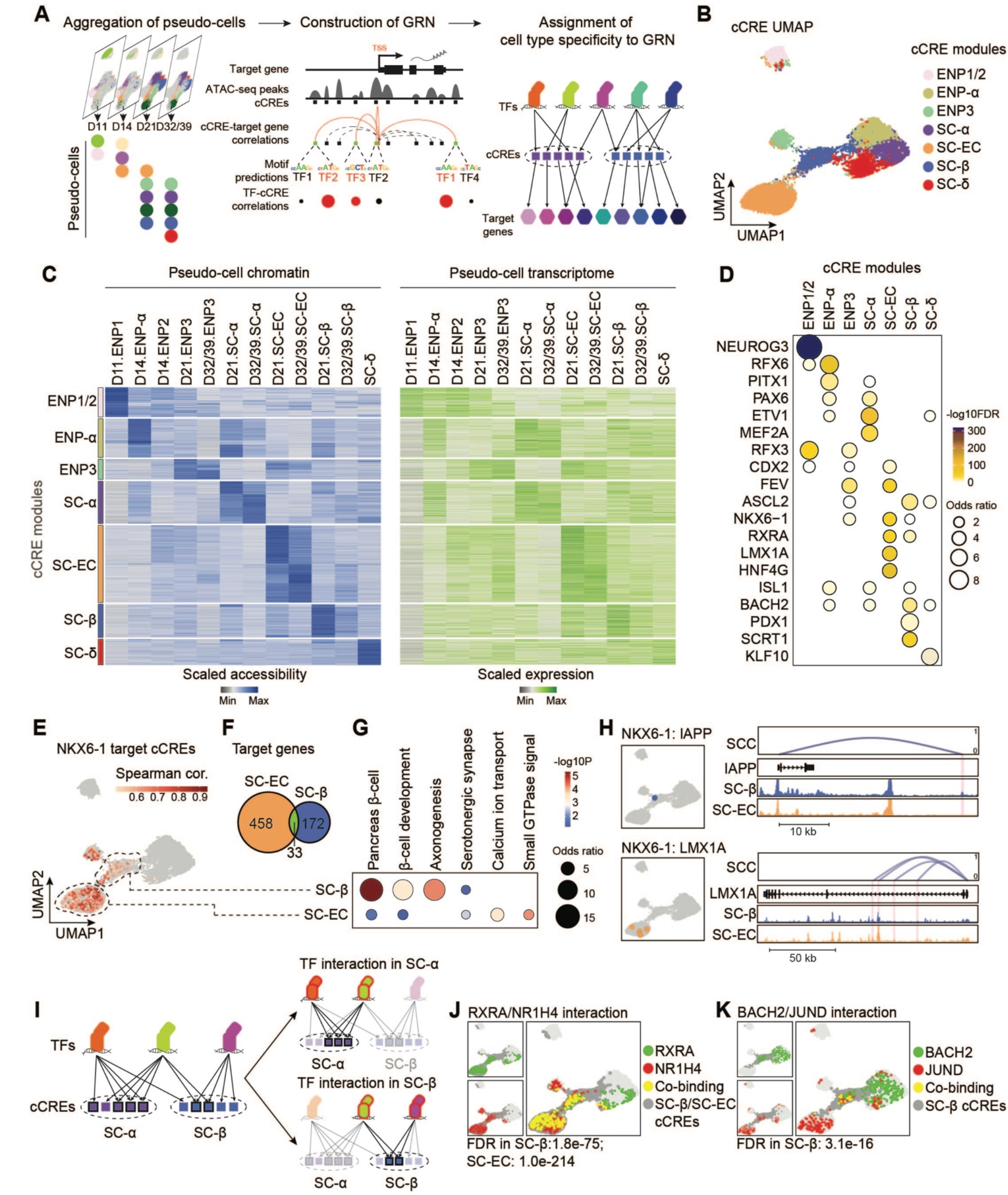
Gene regulatory network analysis of stem cell islet development. (A) Schematic of GRN inference framework and identification of cell type-specific transcriptional programs. Correlation analyses were performed on aggregated pseudo-cells (see Methods). cCRE, candidate *cis* regulatory element. (B) Clustering of GRN cCREs highly variable across cell types and UMAP embedding. Cell identities were assigned to each cCRE module based on cell type with highest chromatin accessibility of the cCREs. (C) Heatmaps showing scaled chromatin accessibility at cCREs (left) and expression levels (right) of target genes linked to the cCRE in each pseudo-cell. (D) Dot plot showing enrichment of TFs predicted to bind to cCREs in each module against a background of all highly variable cCREs. Significance (-log10 FDR) and odds ratio of the enrichments are represented by color and dot size, respectively. (E) UMAP projections of correlations between *NKX6-1* expression and chromatin accessibility of predicted NKX6-1-bound cCREs. SC-β-cell- and SC-EC-specific cCRE modules are highlighted with dashed circles. Spearman cor., spearman correlation coefficient between *NKX6-1* expression and cCRE accessibility. (F) Venn diagram showing overlap between NKX6-1 target genes in SC-ECs and SC-β-cells. Cell type specificity of target genes was determined based on specificity of upstream cCREs. 33 genes are regulated by both SC-β-cell- and SC-EC-specific cCREs. (G) Enriched gene ontology terms/pathways among SC-EC- or SC-β-cell-specific NKX6-1 target genes. Significance (-log10 p-value) and odds ratio of the enrichments are represented by color and dot size, respectively. (H) UMAP locations (left) and genome browser snapshots (right) of predicted NKX6-1-bound cCREs at *IAPP* and *LMX1A* gene loci. Genome browser tracks show aggregated ATAC reads in SC-β-cells and SC-ECs. All tracks are scaled to uniform 1×10^6^ read depth. SCC, spearman correlation coefficients for cCRE accessibility and target gene expression. (I) Schematic of prediction method for cell type-specific TF interactions. (J, K) UMAP projections of predicted TF-TF interactions. Green dots, cCREs bound by background TF; red dots, cCREs bound by test TF; yellow dots, cCREs co-bound by both TFs; dark grey dots, cCRE module(s) with predicted TF interaction(s).

To characterize cell type-specific gene regulatory programs, we subset the GRN by clustering cCREs based on accessibility pattern across cell types (see Methods; **Figure 2B,C** and **Figure S2H**). As expected, cCRE modules specific to related cell types (e.g., ENP-α and SC-α) were localized closely to each other on the cCRE UMAP (**Figure 2B**). Furthermore, target genes linked to cCREs within each module were cell type-specifically expressed (**Figure 2C**) and exhibited cell type-characteristic molecular functions (**Figure S2I** and **Table S3**). Analysis of TFs regulating gene expression in each cell type revealed known regulators of cell type identity, such as NEUROG3 for endocrine progenitors, PAX6 for α-cells, and PDX1 for β-cells (**Figure 2D**, **Figure S2J,** and **Table S4**). Novel predictions included a gene regulatory role for RFX3 and RFX6 in endocrine progenitors, ETV1 in SC-α-cells, ASCL2 and SCRT1 in SC-β-cells, and KLF10 in SC-δ-cells. CDX2, LMX1A, FEV, and HNF4G emerged as candidate regulators of gene expression in SC-ECs and their precursors.

The cell type-specific sub-GRNs allowed us to examine relationships between TFs in the regulation of individual genes as well as target gene specificity of individual TFs across different cell types. For example, we found that cCREs within the *GCG* locus were bound by PAX6 already in ENP-α whereas ETV1 bound to *GCG* cCREs mostly in SC-α-cells (**Figure S2K**), suggesting sequential actions of these TFs in *GCG* gene regulation during α-cell development. NKX6-1 and the retinoic acid X receptor A (RXRA) emerged as candidate regulators of both SC-β-cells and SC-ECs; however, NKX6-1 bound to different cCREs and regulated different genes in SC-β-cells than in SC-ECs (**Figure 2E,F** and **Figure S2L**). Whereas NKX6-1 target genes in SC-β-cells were related to β-cell developmental and β-cell functional processes, NKX6-1 controlled genes involved in serotonergic signaling in SC-ECs (**Figure 2G** and **Table S5**), exemplified by NKX6-1 binding to cCREs in the β-cell gene *IAPP* in SC-β-cells and cCREs in *LMX1A*, a master TF of serotonin synthesis genes (Gross et al., 2016), in SC-ECs (**Figure 2H**). These examples illustrate the power of this analysis for identifying distinct temporal and cell type-specific roles of individual TFs.

We further devised a method to calculate the likelihood of cooperative gene regulation by TFs in a given cell type (see Methods; **Table S6**), inferred based on physical proximity of TF binding within a cCRE and clustering of co-bound cCREs within a cell type-specific cCRE module (**Figure 2I**). We found ENP1-specific cooperativity between the known heterodimers NEUROG3 and TCF3 (Tiedemann et al., 2017) (**Figure S2M**), as well as cooperativity between MAFG and the cap’n’collar (CNC) family TF BACH2 in both SC-α-cells and SC-β-cells (**Figure S2N**), consistent with recruitment of CNC TFs by small MAF proteins (Kannan et al., 2012). RXRA was predicted to cooperate with the bile acid receptor NR1H4 in SC-β-cell and SC-EC gene regulation (**Figure 2J**), identifying a possible mechanism for the role of bile acids in insulin secretion (Vettorazzi et al., 2016). Of interest is also the SC-β-cell-specific interaction between BACH2 and JUND (**Figure 2K**), which have both been reported to contribute to β-cell dysfunction in type 2 diabetes (Good et al., 2019; Son et al., 2021; Wang et al., 2021). In summary, our GRN provides a resource for interrogating gene regulatory mechanisms in SC-islet cell types and their precursors.

### Establishment of SC-islet cell type lineage trajectories

The GRN analysis identified candidate TFs involved in the specification of individual islet cell lineages. However, this analysis left unclear the order in which these TFs function to specify a particular lineage. To gain insight into temporal aspects of islet lineage specification, we ordered gene regulatory programs identified in the GRN along a lineage trajectory by assigning pseudotime values to a given TF, cCREs bound by the TF, and cCRE target genes (see Methods). Using the lineage trajectories for SC-α-cells, SC-β-cells, and SC-ECs (**Figure 1E-H**), we plotted cCRE accessibility across cells for each lineage trajectory (**Figure S3A-C**) and identified maximum cCRE accessibility and target gene expression along each of the three pseudotime lineage trajectories (**Figure S3D-I**). Validating our method, cCREs with early activity in the lineage trajectory - reflected by low cCRE pseudotime values - projected to progenitor-specific cCRE modules in the cCRE UMAP (**Figure 3A,B** and **Figure S3J**).

**Figure 3.**
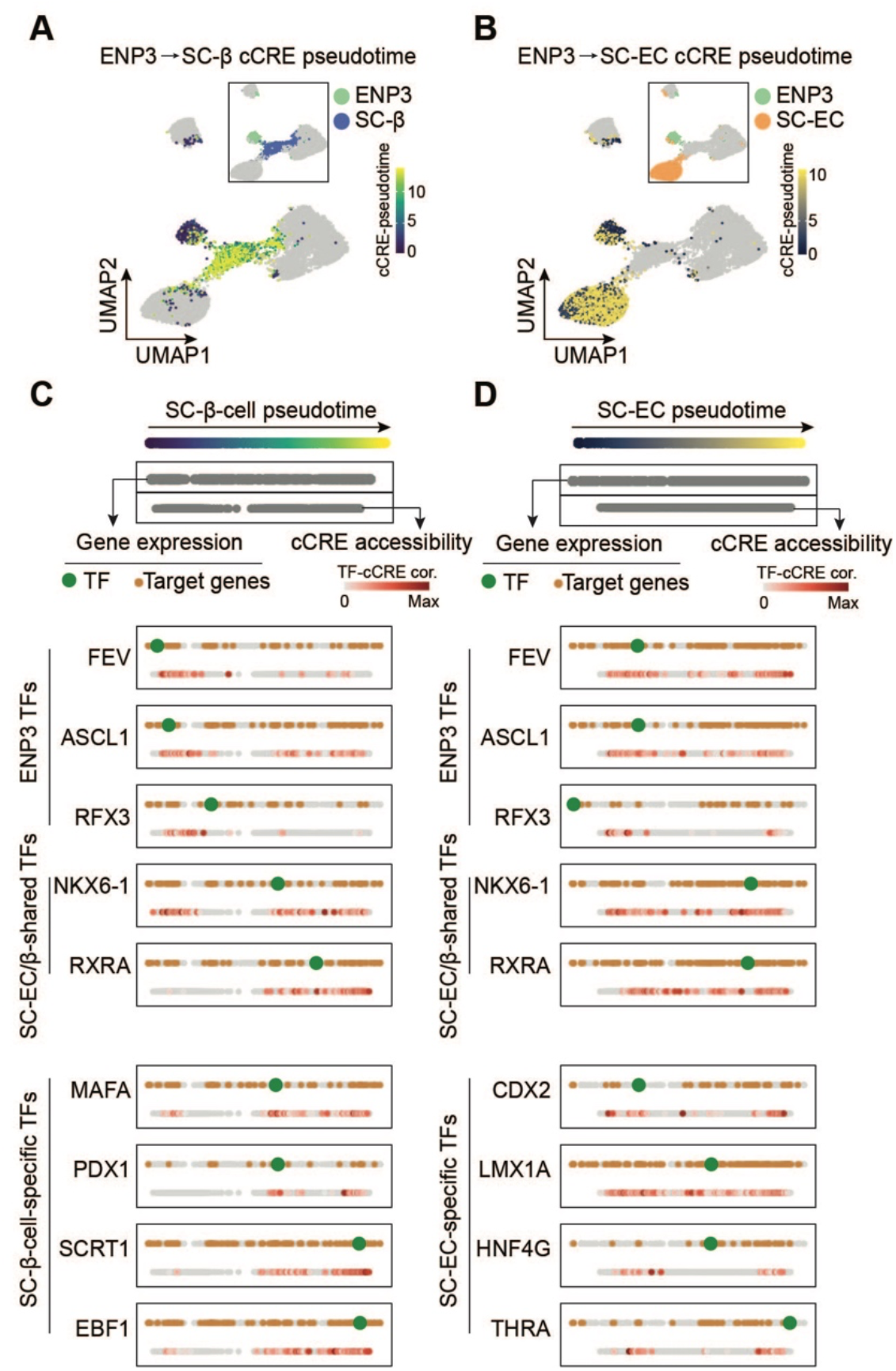
Ordering of transcriptional programs along lineage trajectories. (A, B) UMAP projections of cCRE pseudotime on SC-β-cell (A) and SC-EC (B) lineage trajectories. Insets show cell type annotations of cCRE modules. (C, D) Pseudotime ordering of transcriptional programs along SC-β-cell (C) and SC-EC (D) lineage trajectories from ENP3 progenitors. Gene expression and cCRE accessibility were assigned pseudotime values and plotted in two separate dotted lines (genes, top; cCREs, bottom). For each shown TF, the TF (green), TF-bound cCREs (colored based TF-cCRE correlations) and target genes (brown) are shown.

Integration of both cCRE activity and gene expression into the pseudotemporal lineage trajectory (**Figure 3C,D**, **Figure S3K**, and **Table S7**), allowed us to next quantify the temporal order of TF activity and their downstream target genes during SC-α-cell, SC-β-cell, and SC-EC development. In the SC-α-lineage trajectory, we identified ZNF414, NEUROG3, and PKNOX2 as the earliest TFs, followed by RFX6, PITX1, and PPARG in ENP-α progenitors and ETV1, PAX6, and MEF2C with later functions in SC-α-cell development (**Figure S3K**). While target genes of the early TFs ZNF414 and NEUROG3 remained expressed throughout the trajectory, PKNOX2 target genes were mostly transiently expressed (**Figure S3K**), suggesting distinct gene regulatory programs controlled by these early α-cell lineage TFs. Analysis of the SC-β-cell and SC-EC trajectories from ENP3 progenitors revealed FEV, ASCL1, and RFX3 as ENP3-active TFs with their target genes remaining expressed throughout development of the lineage (**Figure 3C,D**). Later phase TFs in the SC-β-cell and SC-EC trajectories could be separated into two groups: (i) shared TFs between SC-β-cells and SC-ECs, exemplified by NKX6-1 and RXRA and (ii) cell type-specific TFs with MAFA, PDX1, SCRT1, and EBF1 regulating genes in SC-β-cells, and CDX2, LMX1A, HNF4G, and THRA regulating genes in SC-ECs. CDX2 was the earliest TF expressed during SC-EC lineage specification and CDX2-bound cCREs exhibited activity already in ENP3 progenitors (**Figure 3D**), suggesting that CDX2 could have an early role in SC-EC development.

### Identification of an “enterochromaffin cell-like” pre-β-cell population in human fetal pancreas

Since SC-ECs exhibit similarity to enterochromaffin cells of the intestine, they are thought to be islet-aberrant and an erroneous cell type produced in current SC-islet *in vitro* differentiation protocols (Balboa et al., 2022; Veres et al., 2019). Consistent with this view, CDX2 is a well-established master regulator of the intestinal fate (Gao et al., 2009). However, another possibility is that CDX2 is expressed in the developing human pancreas and that SC-ECs represent a *bona fide* pancreatic cell type.

To test the latter possibility, we analyzed scRNA-seq data from 9-19-week post-conception (wpc) fetal human pancreas (Yu et al., 2021) for *CDX2* expression (**Figure S4A**). Indeed, *CDX2* was expressed in ductal/endocrine precursors of the trunk domain as well as in *NEUROG3*^+^/*FEV*^+^ endocrine progenitors (fetal-ENP) and *INS*^+^ fetal β-cells (**Figure S4B**), resembling domains of CDX2 mRNA and protein expression in SC-islets (**Figure S4C,D**). To analyze CDX2 protein in the developing and adult human pancreas, we conducted immunofluorescence staining on fetal (10, 13, 18, and 20 wpc), neonatal (1 day, 2 months, and 3.7 months postnatally), and adult pancreas (**Figure 4A-D**). In both fetal and neonatal pancreas, CDX2 was expressed in PDX1^+^ pancreatic progenitors and in a small subset of β-cells. In adult pancreas, CDX2 was found in ductal structures outside the islets but was no longer detected in β-cells. Thus, CDX2 is expressed in endocrine progenitors and a subset of β-cells of the developing human pancreas, suggesting that SC-ECs could present a transient, fetal β-cell precursor population.

**Figure 4.**
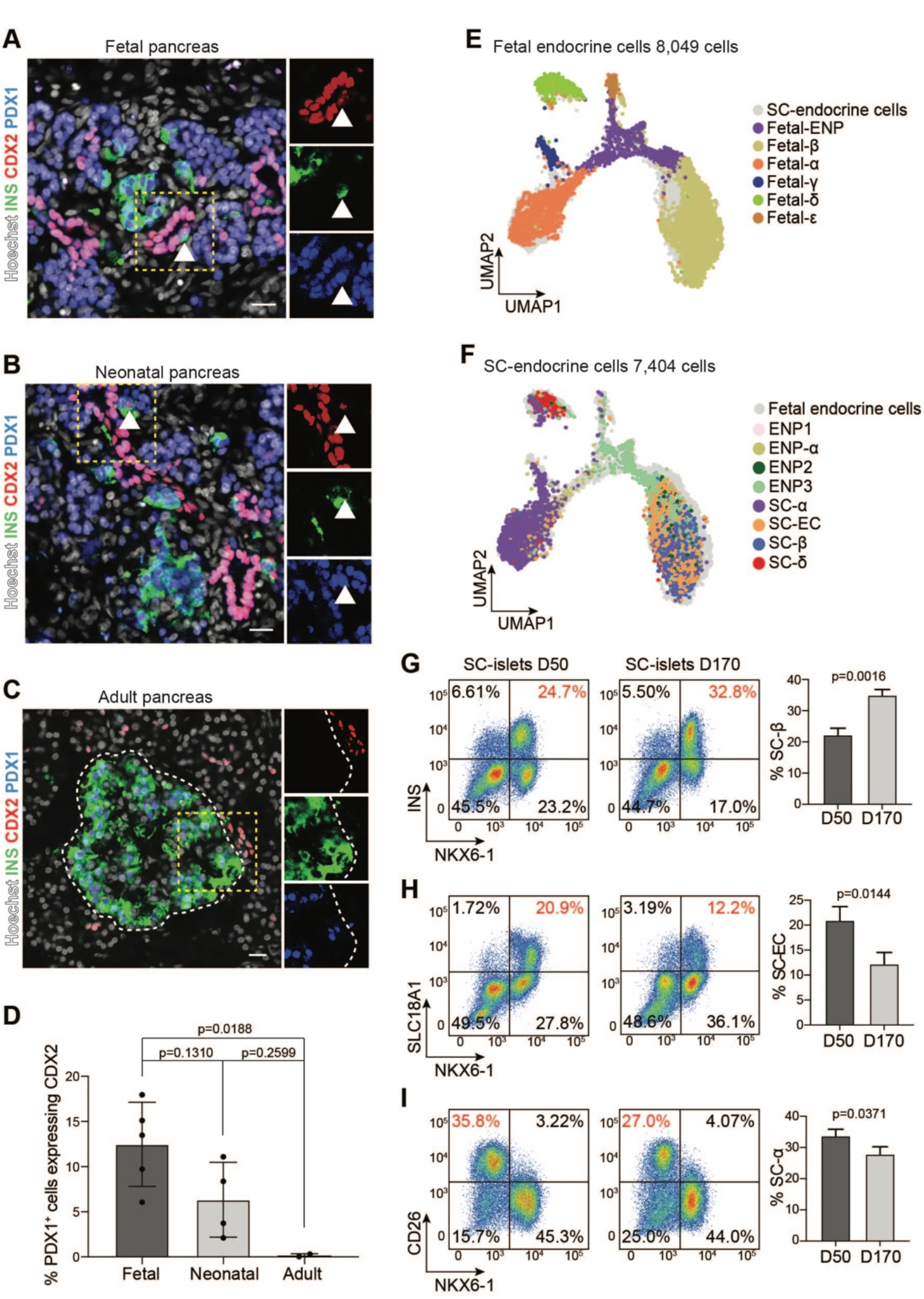
A transient, CDX2^+^ pre-β-cell population in fetal pancreas resembles stem cell-derived enterochromaffin cells. (A-C) Representative immunofluorescent images for CDX2, PDX1, and insulin (INS) on human pancreas at 10 weeks post conception (wpc, A), postnatal day 1 (B), and 52 years of age (C). Arrowheads in insets indicate CDX2 and INS co-positive cells. Nuclei were labeled with Hoechst. Scale bar, 20 μm. (D) Quantification of PDX1^+^ cells expressing CDX2 in fetal (10-20 wpc, n = 5), neonatal (1 day to 3.7 months postnatally, n = 4), and adult (20-52 years, n = 2) human pancreas. Data are shown as mean ± S.D. P-values were calculated using Tukey’s multiple comparisons test after one-way ANOVA. (E, F) UMAP co-embedding of single-cell transcriptomes from endocrine cells in fetal human pancreas (E) and during SC-islet differentiation (F). Cells are color-coded based on their annotated identities in Extended Data Figure 4a and Figure 1b, respectively. (G-I) Representative flow cytometry plots (left, percentage of population of interest in red) and quantifications (right) of SC-β-cells (NKX6-1^+^/INS^+^, G), SC-ECs (NKX6-1^+^/SLC18A1^+^, H) and SC-α-cells (NKX6-1^-^/CD26^+^, I) in early (day (D) 50) and late (D170) SC-islet cultures. Data are shown as mean ± S.D. (n = 3 independent differentiations). P-values were calculated by unpaired two-tailed t-test.

To assess the transcriptomic similarity between primary fetal β-cells and SC-ECs, we co-embedded scRNA-seq data from human fetal endocrine cells (**Figure S4A**) and cells during SC-islet differentiation (**Figure 1B**) on the same UMAP, allowing for similarity assessment of cell types based on proximity on the UMAP. Interestingly, both SC-ECs and SC-β-cells co-localized with fetal β-cells (**Figure 4E,F**), demonstrating that SC-β-cells and SC-ECs both represent a fetal β-cell state and that SC-ECs are not pancreas-aberrant. Since an SC-EC-like population is no longer present in adult pancreas (**Figure 4C,D** and (Veres et al., 2019)), we determined whether SC-ECs decrease in numbers as SC-islets functionally mature. Long-term SC-islet culture induced β-cell functional maturation (**Figure S4E**), akin to β-cell maturation characteristic of postnatal development (Aguayo-Mazzucato et al., 2011; Arda et al., 2016; Henquin and Nenquin, 2018; Jermendy et al., 2011; Otonkoski et al., 1988; Rorsman et al., 1989). Compared to short-term cultures (D50), long-term SC-islet cultures (D170) comprised a higher percentage of SC-β-cells (NKX6-1^+^/INS^+^; **Figure 4G**) and lower percentage of SC-ECs (NKX6-1^+^/SLC18A1^+^; **Figure 4H**), whereas the percentage of SC-α-cells (NKX6-1^-^/CD26^+^(Molakandov et al., 2021)) was unchanged (**Figure 4I**), supporting that SC-ECs are a transient, β-cell-related population. Collectively, our findings identify a novel CDX2^+^ β-cell precursor-like population in the human fetal pancreas with close resemblance to SC-ECs produced during SC-islet cell differentiation.

### CDX2 regulates serotonin synthesis genes

The GRN analysis suggests that CDX2 is a critical TF for gene regulation in endocrine progenitors and SC-ECs (**Figure 2D**). Examination of CDX2 target genes revealed several genes involved in serotonin synthesis, including tryptophan hydroxylase (*TPH1*) and the serotonin transporter *SLC18A1*. Both genes are expressed in endocrine progenitors and β-cells of the fetal pancreas as well as SC-derived ENP3 progenitors and SC-ECs (**Figure 5A**). At the *TPH1* locus, CDX2 bound to a distal cCRE highly active in SC-ECs and fetal endocrine cells (Domcke et al., 2020) but not SC-β-cells or fetal pre-ductal/endocrine and pre-acinar cells (**Figure 5B** and **Figure S5A**). A similar chromatin activity pattern was found at the CDX2-bound *SLC18A1* promoter (**Figure 5C** and **Figure S5B**). Thus, a program of serotonin synthesis genes is expressed in a subset of endocrine progenitors and β-cells of the human fetal pancreas and this program is predicted to be regulated by CDX2.

**Figure 5.**
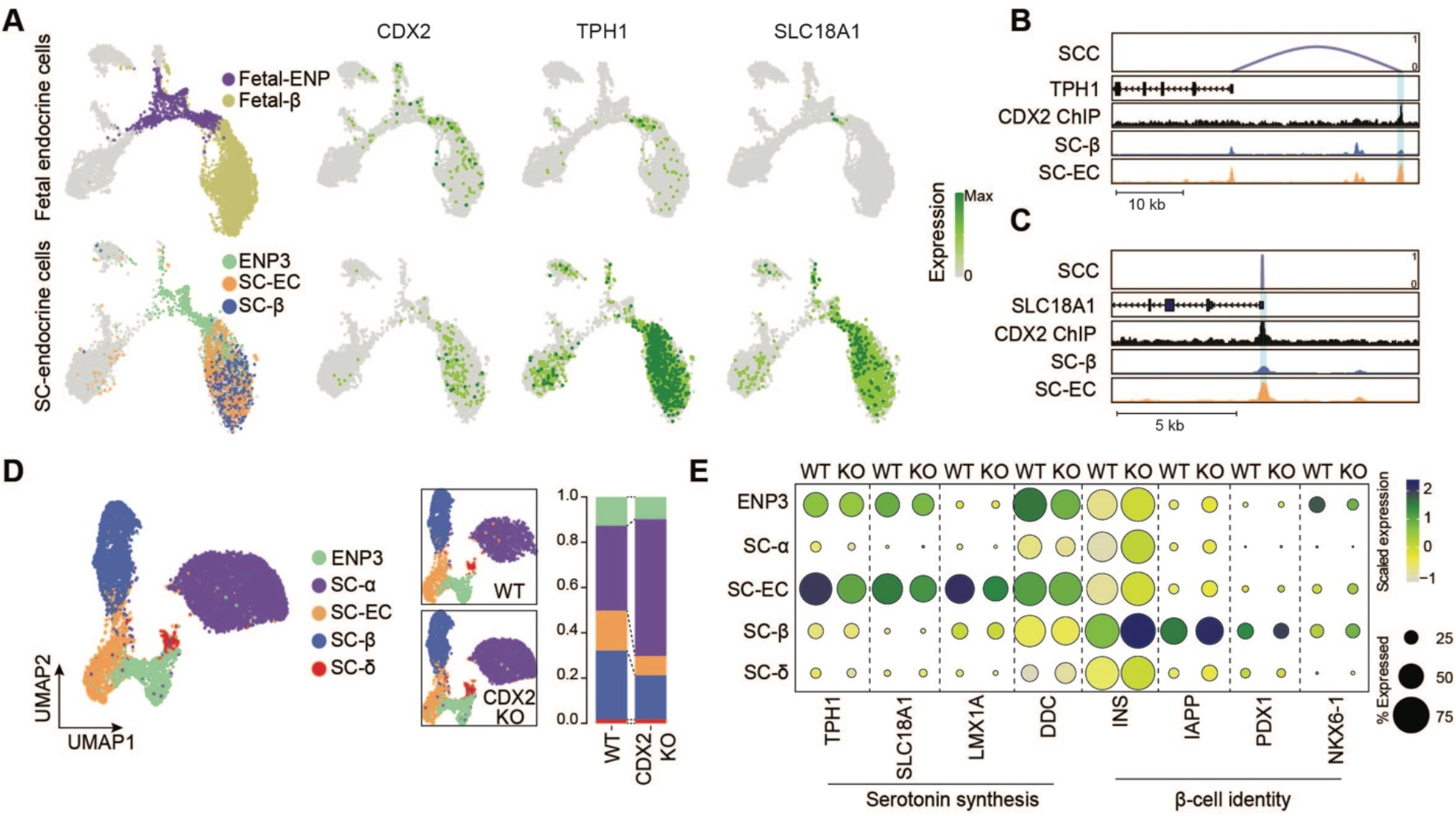
CDX2 regulates serotonin synthesis genes. (A) Expression of *CDX2*, *TPH1,* and *SLC18A1* in fetal (top) or stem cell-derived (bottom) endocrine cells. UMAPs on left indicate location of relevant cell types. (B, C) Genome browser tracks showing CDX2 ChIP-seq reads in SC-islets and aggregated ATAC reads in SC-β-cells and SC-ECs at *TPH1* (B) and *SLC18A1* (C) gene loci. CDX2-bound cCREs are highlighted. All tracks are scaled to uniform 1×10^6^ read depth. SCC, spearman correlation coefficients for cCRE accessibility and target gene expression. (D) UMAP co-embedding of single cell transcriptomes from wild type (WT) and *CDX2* knockout (KO) SC-islets. Cells are color-coded based by transferred identities from Figure 1b. The relative abundance of each cell type in WT and *CDX2* KO SC-islets is shown on the right. (E) Dot plot showing differentially expressed genes in WT and *CDX2* KO SC-islet cell types. The color of each dot represents the expression level and the size the percentage of cells expressing the gene.

To examine CDX2 function during islet cell development, we deleted *CDX2* in hESCs (*CDX2*-KO line) by introducing a frameshift and premature termination codon into the *CDX2* coding region using CRISPR/Cas9-mediated genome editing (**Figure S5C-E**). We differentiated *CDX2*-KO hESCs and unedited wildtype (WT) hESCs into SC-islets and quantified endocrine cell type composition by flow cytometry and image analysis. *CDX2* inactivation led to a significant reduction in SC-ECs, a slight reduction in SC-β-cells, but no significant change in SC-α-cells (**Figure S5F-K**). Single-cell RNA-seq analysis confirmed the reduction in SC-ECs in *CDX2*-KO compared to WT SC-islets and revealed a decrease in SC-β-cells and increase in SC-α-cells (**Figure 5D**), suggesting that CDX2 controls the lineage decisions at the branchpoint between SC-α-cells and SC-ECs/β-cells. Of note, the discrepancy between effects of *CDX2* deletion on cell population composition by marker protein analysis and scRNA-seq is likely explained by marker proteins only capturing a small aspect of cell identity. The scRNA-seq analysis further revealed a significant reduction in the expression of serotonin synthesis genes (*TPH1*, *SLC18A1*, *LMX1A*, *DDC*) in *CDX2*-KO ENP3 or SC-ECs, whereas genes characteristic of β-cell identity (*INS*, *IAPP*, *PDX1*, *NKX6-1*) were more highly expressed in SC-ECs and SC-β-cells after *CDX2* deletion (**Figure 5E, Figure S5L**, and **Table S8**). These findings suggest that CDX2 expression favors adoption of SC-EC over SC-β-cell identity.

Together, our analysis suggests that CDX2 is a marker of a transient *bona fide* fetal pancreatic cell population with pre-β-cell identity. We show that CDX2 drives the expression of serotonin synthesis genes in these pre-β-cells. Serotonin production is a feature of neonatal and adolescent β-cells (Castell et al., 2022; Moon et al., 2020b), and adult β-cells can activate serotonin synthesis during pregnancy (Baeyens et al., 2016; Kim et al., 2010; Moon et al., 2020a; Ohara-Imaizumi et al., 2013). Based on this evidence, we posit that SC-ECs are not an erroneous intestinal cell type of SC-islet *in vitro* differentiation but rather represent a cell type characteristic of normal human β-cell development.

### Insufficient activation of signal-dependent gene regulatory programs in SC-islets

Having established gene regulatory programs for each SC-islet cell type, we sought to determine how closely these programs resemble those of corresponding islet cell types in postnatal pancreas. We reasoned that this knowledge could inform strategies to further mature SC-β-cells *in vitro*. Toward this goal, we generated snATAC-seq, scRNA-seq, and single-nucleus RNA-sequencing (snRNA-seq) datasets from primary islets and pancreas from childhood (ages 13 months to 9 years) and adult donors (ages 20-66) and complemented these data with publicly available islet scRNA-seq data **(Table S1A,B**).

To focus the analysis on endocrine cell types, we selected endocrine populations from each dataset for data integration (SC populations: ENP3, ENP-α, SC-β, SC-α, SC-δ, and SC-EC; primary cell populations: α-, β-, δ-, and γ-cells; **Figure 6A** and **Figure S6A-H**). After batch correction of scRNA-seq and snRNA-seq data (**Figure S6C,D,G,H**), we integrated data from SC-derived and primary cells into one UMAP for chromatin accessibility and gene expression, respectively (**Figure 6B,C**). For both chromatin accessibility and gene expression, we identified a single α-cell, δ-cell, and γ-cell cluster as well as a population of β-cells comprised of four subclusters (**Figure 6B,C** and **Figure S6I**). Chromatin accessibility and gene expression data were highly concordant between clusters (**Figure S6J**) and cell type annotations in the integrated map largely corresponded to cell identities prior to data integration (**Figure S6K,L**). The α-cell and δ-cell clusters each comprised α-cells and δ-cells from stem cells, childhood, and adult pancreas, demonstrating a high degree of similarity between SC-α- and SC-δ-cells with their primary counterparts (**Figure 6B,C** and **Figure S6K,L**). Of note, a subset of SC-α-cells clustered with primary γ-cells in the integrated map (**Figure 6B,C**, dashed circles), consistent with developmental similarity between α-cells and γ-cells (Baron et al., 2016). In contrast to non-β-cell endocrine cell types, SC-β-cells clustered separately from primary β-cells. While the majority of primary β-cells from childhood and adult donors clustered together (cluster β-1), SC-β-cells, SC-ECs, and ENP3 progenitors each localized to a distinct β-cell cluster (clusters β-2, β-3, and β-ENP). This suggests that SC-β-cells are more distant to their primary counterparts than other SC-derived endocrine cell types.

**Figure 6.**
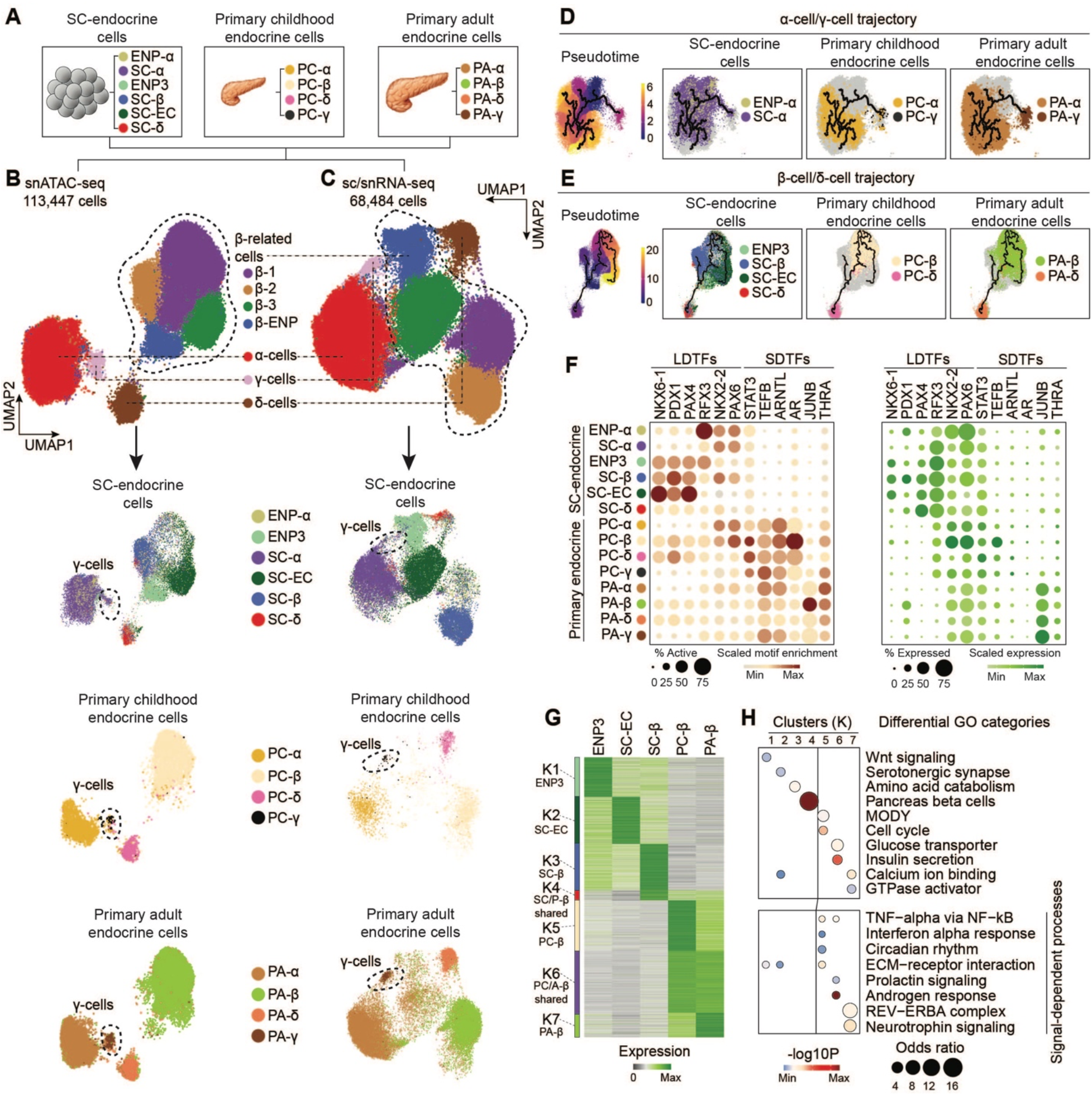
Insufficient activation of signal-dependent gene regulatory programs in stem cell β-cells. (A) Schematic showing cell types included into the integrative analysis of snATAC-seq and sc/snRNA-seq data. Stem cell (SC)-derived endocrine cell types and endocrine progenitors (SC-α-cells, SC-β-cells, SC-ECs, SC-δ-cells, ENP-α, and ENP3), primary endocrine cell types from childhood pancreas (PC-α-cells, PC-β-cells, PC-δ-cells, and PC-γ-cells) and primary endocrine cell types from adult pancreas (PA-α-cells, PA-β-cells, PA-δ-cells, and PA-γ-cells) were included. (B, C) UMAP embedding of chromatin accessibility (B) and transcriptome (C) data from cell types detailed in (A). Cluster identities were defined by promoter accessibility (snATAC-seq) or expression (sc/snRNA-seq) of marker genes. The dashed line outlines β-cell-related cell types. Bottom panels: split UMAPs showing localization of stem cell, childhood and adult pancreatic endocrine cells. Cells were color-coded based on their identities from (A). (D, E) Trajectory analysis based on chromatin accessibility, showing trajectories for α-cells/γ-cells (D) and β-cells/δ-cells (E) with ENP-α and ENP3 set as the root, respectively. Cells were color-coded by either original identities (A) or pseudotime values. (F) Dot plots showing scaled average motif enrichment (left) or gene expression (right) of TFs. The color of each dot represents the average motif enrichment or expression level and the size of each dot the percentage of positive cells for each TF. LDTF, lineage-determining TF; SDTF, signal-dependent TF. (G) K-means clustering of genes with variable expression across β-related cell types (ENP3, SC-ECs, SC-β-cells, PC-β-cells, and PA-β-cells). Clusters were annotated and color-coded based on gene expression patterns. (H) Enriched gene ontology terms/pathways in each cluster. Significance (-log10 p-value) and odds ratio of the enrichments are represented by color and dot size, respectively.

We sought to further analyze the relatedness of SC-islet cells to primary endocrine cells by inferring lineage trajectories based on chromatin accessibility. We built two separate trajectories by grouping cell types with known lineage relationship (Baron et al., 2016): (i) an α-cell/γ-cell trajectory with ENP-α, SC-α-cells, and primary childhood and adult α- and γ-cells (**Figure 5D**); and (ii) a β-cell/δ-cell trajectory with ENP3, SC-EC, SC-β-cells, SC-δ-cells, and primary childhood and adult β- and δ-cells (**Figure 6E**). In the α-cell/γ-cell trajectory, we observed a progression from ENP-α to SC-α-cells to childhood α-cells, and finally to adult α-cells, suggesting that SC-α-cells are less mature than primary α-cells but on the correct path. A subset of SC-α-cells localized on a separate branch together with primary γ-cells, indicating γ-cell lineage potential of this subset of SC-α-cells. In the β-cell/δ-cell trajectory, we observed three trajectories each originating from ENP3 progenitors. One branch encompassed SC-δ-cells and primary δ-cells, one SC-β-cells, and a third primary β-cells with SC-ECs being closely associated. This analysis confirms the relatedness of SC-ECs to β-cells, providing further support for the notion that SC-ECs represent a unique β-cell state in human development.

To identify gene regulatory programs that distinguish primary from SC-derived endocrine cell types, we identified TF motifs with variable enrichment in accessible chromatin across cell populations (see Methods; **Figure 6F** and **Table S9**). Motifs for lineage-determining TFs, such as NKX6-1, PDX1, NKX2-2, and PAX6, were enriched in SC-islet cell types, suggesting that gene regulatory programs driven by lineage-determining TFs are sufficiently active in SC-islet cells. By contrast, we observed enrichment for signal-dependent TF motifs in primary compared to SC-islet endocrine cell populations, consistent with lower expression of some of these TFs in SC-islet cells (**Figure 6F**). These signal-dependent TFs included STAT3 which is activated by signals from immune cells (De Groef et al., 2016; Linnemann et al., 2017), the circadian clock-dependent TF ARNTL (Alvarez-Dominguez et al., 2020; Perelis et al., 2015; Rakshit et al., 2018; Vieira et al., 2012), and TFs activated by steroid hormones, such as the androgen receptor (AR) and thyroid hormone receptor (THRA) (Aguayo-Mazzucato et al., 2013; Bruin et al., 2015; Rezania et al., 2014). These findings suggest that insufficient activation of signal-dependent gene regulatory programs distinguishes SC-derived from primary endocrine cells.

To next define differences in gene expression between SC-islet and primary endocrine cell types, we identified differentially expressed genes between β-lineage cell types (ENP3, SC-β-cells, SC-ECs versus primary childhood β-cells versus adult β-cells) and α-lineage cell types (SC-α-cells versus primary childhood α-cells versus adult α-cells), respectively, by pairwise differential gene expression analysis. Clustering of differentially expressed genes identified gene modules that distinguish stem cell-derived from primary cells as well as shared modules (**Figure 6G** and **Figure S6M**). Genes in modules more highly expressed in primary β-cells than SC-β-related cells were associated with signaling pathways known to regulate β-cell function, including inflammatory, circadian, and neurotrophin signaling (**Figure 6H** and **Table S10**). A similar lack of activation of signal-dependent gene expression was observed in SC-α-cells compared to primary α-cells (**Figure S6N** and **Table S11**). In addition, genes associated with insulin secretion were expressed at higher levels in primary β-cells, whereas SC-β-cells expressed higher levels of genes related to amino acid catabolism (**Figure 6H**), consistent with a more pronounced insulin secretory response to amino acids in SC-β-cells compared to primary β-cells (Balboa et al., 2022; Helman et al., 2020). Together, these findings underscore that important signaling events are not sufficiently induced in SC-derived endocrine cells and suggest that insufficient activation of these signal-dependent processes could account for remaining functional differences between primary and SC-β-cells.

### Steroid hormones stimulate β-cell proliferation

To comprehensively identify TFs and associated target genes with differential activity in primary and SC-β-cells, we constructed a GRN comprised of β-cell-related populations analogous to the approach described in Figure 2a (see Methods). We connected cCREs to their target genes (FDR<0.15; **Figure S7A-C**) and cCREs to upstream TFs (Spearman correlation>0 and FDR<0.05; **Figure S7F**), identifying a total of 377 TFs connected to 96,020 cCREs and 12,370 target genes. Each TF in the network bound an average of 2,698 cCREs, each cCRE regulated 1.55 genes, and each gene was regulated by 13.6 cCREs (**Figure S7D,E,G**).

We then identified cCREs in the GRN with highly variable chromatin accessibility across cell populations, clustered cCREs of β-cell populations (ENP3, SC-EC, SC-β, and primary childhood and adult β-cells), and plotted cCRE modules onto a UMAP (see Methods). In addition to population-specific modules, we identified cCRE modules shared between cell types, exemplified by a shared module between childhood and adult primary β-cells, SC-β-cells and primary β-cells, and SC-ECs and primary β-cells (**Figure 7A** and **Figure S7H**). The shared SC-β-cell/primary β-cell module lies between the SC-β-cell- and childhood β-cell-specific modules, suggesting that aspects of gene regulatory changes associated with β-cell maturation occur in SC-β-cells. Furthermore, presence of a SC-EC/primary β-cell module indicates that SC-ECs share some gene regulatory features with primary β-cells, supporting their relatedness.

**Figure 7.**
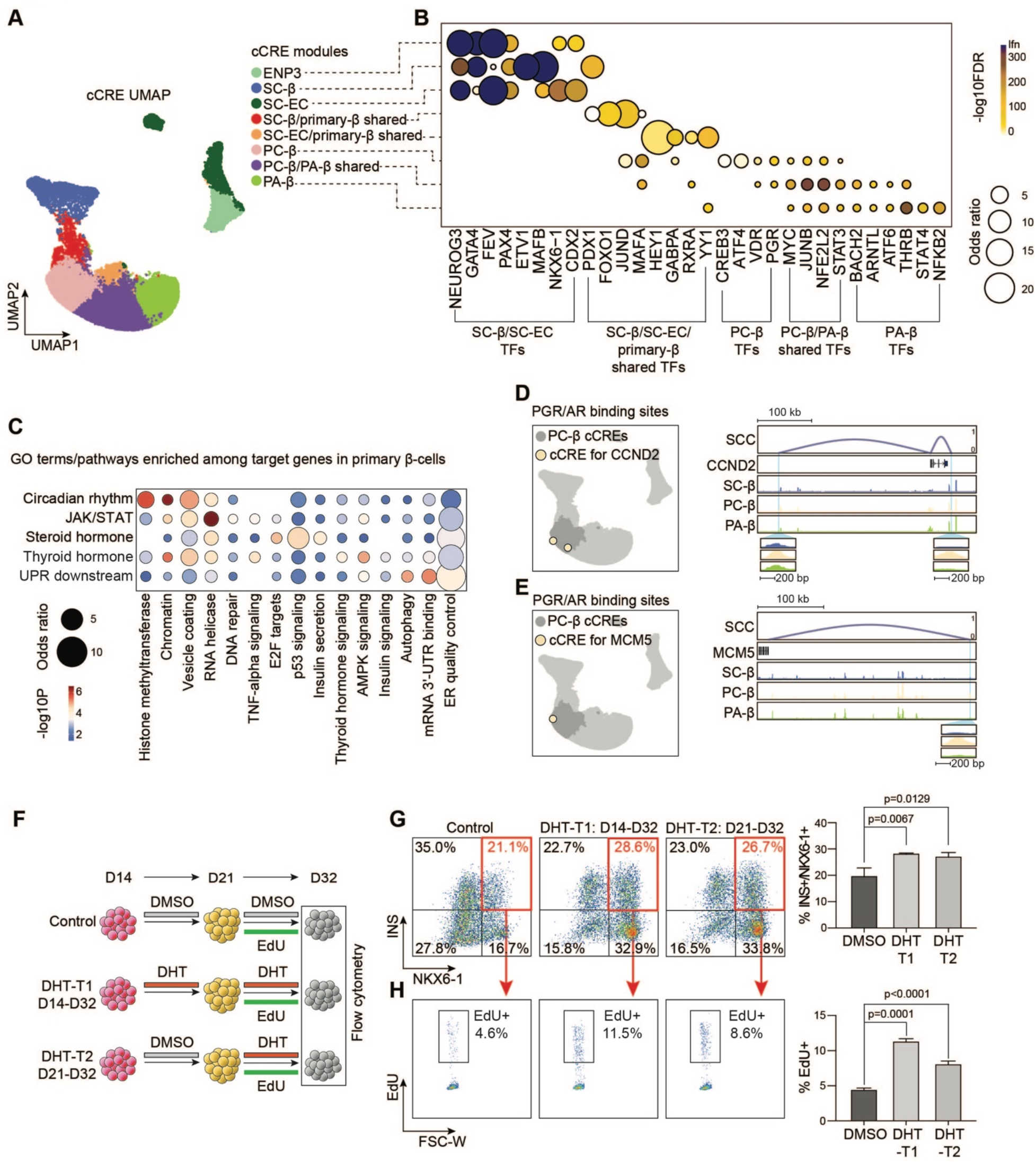
Gene regulatory network underlying β-cell maturation. (A) Clustering of GRN cCREs highly variable across β-related cell types and UMAP embedding. Cell identities were assigned to each cCRE module based on cell type with highest chromatin accessibility of the cCREs. ENP, endocrine progenitor; SC, stem cell; EC, enterochromaffin-like cell; PC, primary childhood; PA, primary adult. (B) Dot plot showing enrichment of TFs predicted to bind to cCREs in each module. Significance (-log10 FDR) and odds ratio of the enrichments are represented by color and dot size, respectively. (C) Enriched gene ontology terms/pathways among target genes regulated by signals active in primary β-cells (PC-β and PA-β combined). Significance (-log10 p-value) and odds ratio of the enrichments are represented by color and dot size, respectively. (D, E) UMAP locations (left) and genome browser snapshots (right) of predicted PGR/AR-bound cCREs at *CCND2* (D) and *MCM5* (E) gene loci. Genome browser tracks show aggregated ATAC reads in SC-β-cells, PC-β-cells, and PA-β-cells. PGR/AR-bound PC-β-cell-specific cCREs at *CCND2* (D) and *MCM5* (E) are highlighted. All tracks are scaled to uniform 1×10^6^ read depth. SCC, spearman correlation coefficients for cCRE accessibility and target gene expression. (F) Experimental design for dihydrotestosterone (DHT) treatment of SC-islets. SC-islets were treated with 10 nM DHT or DMSO from D14-D32 or D21-D32. The nucleoside analogue EdU was added at 10 µM to label proliferating cells. (G) Representative flow cytometry plots (left, SC-β-cell percentage in red) and quantifications (right) of SC-β-cells (NKX6-1^+^/INS^+^) in D32 SC-islets with treatments shown in (F). Data are shown as mean ± S.D. (n = 3 independent differentiations). P-values were calculated by Dunnett’s multiple comparisons test after one-way ANOVA. (H) Representative flow cytometry plots (left) and quantifications (right) of EdU^+^ SC-β-cells (NKX6-1^+^/INS^+^) in D32 SC-islets with treatments shown in (F). Data are shown as mean ± S.D. (n = 3 independent differentiations). P-values were calculated by Dunnett’s multiple comparisons test after one-way ANOVA.

Analysis of TFs with different activity across modules confirmed activation of programs downstream of lineage-determining TFs (e.g., FEV, PAX4, NKX6-1, CDX2, PDX1) and insufficient activation of programs regulated by signal-dependent TFs (e.g., PGR, VDR, STAT3, ARNTL, ATF6, THRA/B) in SC-derived β-cell populations (**Figure 7B** and **Table S12**).

To comprehensively catalog signal-dependent molecular processes insufficiently activated in SC-β-cells, we grouped signal-dependent TFs exclusively active in primary β-cells based on upstream signals regulating their activity and identified downstream target genes from the GRN. Primary β-cell-specific signaling pathways included circadian rhythm (ARNTL, NPAS2), interleukins (STAT1-4, STAT5A, STAT5B, STAT6), steroid hormones (AR, PGR, ESR1, ESR2), thyroid hormones (THRA, THRB), and the unfolded protein response (UPR; ATF6, ATF6B, ATF4) (**Figure 7C** and **Table S13**). Validating the approach, genes downstream of the thyroid hormone receptor were involved in thyroid hormone signaling and TFs of the UPR regulated genes involved in endoplasmic reticulum (ER) quality control (**Figure 7C** and **Table S13B,D**). The analysis predicted regulation of chromatin modifiers by circadian cues, suggesting a role for circadian signals in modulating the β-cell epigenome. Furthermore, identification of thyroid hormone as an upstream regulator of genes involved in AMPK signaling established a molecular link between thyroid hormone and AMPK signaling, which have been independently identified as inducers of β-cell maturation (Aguayo-Mazzucato et al., 2013; Helman et al., 2020; Jaafar et al., 2019). TFs activated by the UPR were identified as regulators of autophagy genes in β-cells, consistent with experimental data showing a role for autophagy in the regulation of β-cell function under ER stress (Bachar-Wikstrom et al., 2013; Bugliani et al., 2019). Together, the GRN provides a framework for understanding signal-dependent regulation of molecular processes in β-cells and identifies signal-dependent processes insufficiently activated during *in vitro* β-cell differentiation.

A striking finding from the network analysis was that steroid hormones were predicted to regulate E2F targets (**Figure 7C** and **Table S13C**), which suggests a role for steroid hormone signaling in β-cell proliferation. The childhood β-cell-specific module was enriched for genes regulated by the progesterone receptor (PGR) which shares a sequence motif with the androgen receptor (AR). We validated PGR/AR motif enrichment in childhood β-cells in H3K27ac ChIP-seq data from sorted childhood compared to adult human β-cells (**Figure S7I**). This finding indicates that sex hormones could promote β-cell proliferation specifically during the childhood period. Accordingly, the GRN revealed the cell cycle genes *CCND2* and *MCM5* as targets of PGR/AR signaling in the childhood β-cell-specific cCRE module (**Figure 7D,E**). Given that sex hormone-driven expression of cell cycle genes was mostly enriched in primary childhood β-cells (**Figure 7B** and **Table S12**), we postulated that activation of the PGR or AR receptor could induce SC-β-cell proliferation. To test this hypothesis, we treated SC-islets with dihydrotestosterone (DHT) during two different time windows of SC-islet differentiation and quantified relative β-cell numbers and proliferation rates (**Figure 7F**). During both treatment windows, DHT significantly increased SC-β-cell numbers and proliferation assessed by EdU incorporation (**Figure 7G,H**). These results identify a previously unknown role for AR signaling in β-cell proliferation, which suggests a possible connection between the surge in neonatal testosterone (Clarkson and Herbison, 2016; Corbier et al., 1992) and early postnatal β-cell proliferation.

## Discussion

It is still a major challenge to influence cell fate decisions during SC-islet differentiation and a roadmap for maturing *in vitro*-produced β-cells is missing. Here we integrated transcriptome and chromatin accessibility data from SC-islets and primary islets to gain insight into the mechanisms underlying cell fate choices during SC-islet differentiation as well as to benchmark gene regulatory programs of SC-islets against those of primary islets. From the integrated data we inferred GRNs and cell type-specific gene regulatory modules, which define regulatory relationships between TFs, TF-binding sites, and target gene expression levels in each given cell type. In addition to identifying novel transcriptional regulators of each SC-islet cell type (i.e., PITX1 and ETV1 for SC-α-cells, SCRT1 and EBF1 for SC-β-cells, and CDX2 for SC-ECs), our integrated GRN provides a global framework for understanding gene regulatory mechanisms of islet cell fate acquisition and maturation. The here-defined GRNs predict the temporal order in which TFs function to specify each lineage and resolve mechanisms of synergy between TFs with temporal and spatial resolution. This information provides a rich resource to design experiments for programming specific islet cell types and maturation states.

Previous work has described production of serotonin-producing cells during SC-islet differentiation (Balboa et al., 2022; Veres et al., 2019). Based on their similarity to intestinal enterochromaffin cells and absence of a similar cell type in adult islets, it has been proposed that these cells are pancreas-aberrant and lack a direct lineage relationship with β-cells (Veres et al., 2019). Our findings show that SC-ECs have close similarity to a CDX2-expressing, serotonin-producing cell population in the fetal pancreas. These cells exhibit endocrine progenitor identify and a subset expresses insulin, suggesting that these cells mark a transition state from endocrine progenitor to early β-cell. Therefore, SC-ECs may represent a β-cell precursor during SC-islet development supported by observations that SC-ECs decrease in abundance during *in vitro* maturation (**Figure 4H**) and after SC-islet engraftment (Balboa et al., 2022). Whether the identified fetal pre-β-cell population represents a transitory state through which all progenitors progress during β-cell differentiation or whether only a subset of adult β-cells arise from this population is still unclear. We observed that a subset of primary β-cells in adult pancreas shares epigenomic features with SC-ECs (**Figure S6K**), which could indicate a distinct developmental origin of this β-cell subset. However, additional studies will be necessary to determine to which extent adult β-cell heterogeneity is developmentally determined. An interesting question in this context is whether the observed reactivation of the fetal serotonin synthesis program during pregnancy (Kim et al., 2010) is restricted to a subpopulation of β-cells or occurs in all β-cells. Regardless, given that SC-ECs are part of the islet lineage, proposed SC-EC depletion strategies (Veres et al., 2019) might not be necessary for a SC-islet cell therapy.

Whereas β-cell differentiation occurs prenatally, the neonatal and early childhood period is characterized by the expansion, proliferation, and functional maturation of β-cells (Aguayo-Mazzucato et al., 2011; Arda et al., 2016; Henquin and Nenquin, 2018; Jermendy et al., 2011; Otonkoski et al., 1988; Rorsman et al., 1989). Postnatal changes in β-cells are thought to be driven by environmental cues (Wortham and Sander, 2021); however, the specific signals have remained poorly characterized. We found that lineage-determining TFs engage with the genome in SC-β-cells, indicating proper differentiation *in vitro*. By contrast, there was a profound lack of engagement of signal-dependent TFs in SC-β-cells compared to primary β-cells. Our integrated GRN identified insufficient activation of circadian, JAK/STAT, steroid and thyroid hormone, as well as UPR signals in SC-β-cells. While circadian cues and thyroid hormone are known β-cell maturation signals (Aguayo-Mazzucato et al., 2013; Rakshit et al., 2018), the specific roles of the other signals remain to be studied. The GRN indicates that JAK/STAT-mediated regulation of stress response genes distinguishes primary from SC-β-cells, which could be due to the absence of islet-resident immune cells in SC-islets. Of interest is the predicted link between UPR signaling and autophagy in β-cells. Experimental data support regulation of autophagy by the UPR (Bachar-Wikstrom et al., 2013; Bugliani et al., 2019), but it remains to be explored whether the UPR and autophagy play a role in human β-cell maturation.

From our analysis, sex hormone-mediated activation of proliferation genes emerged as a gene regulatory program specific to childhood β-cells. We experimentally confirmed this prediction, demonstrating accuracy of the GRN for predicting gene regulatory processes in human β-cells. Interestingly, a similar pro-proliferative effect of androgens has been reported during neurogenesis in human brain organoids (Kelava et al., 2022), suggesting a shared mechanism between pancreatic and neuronal cells. Our findings predict a role for sex hormones in postnatal β-cell expansion. Stimulation of sex hormone-dependent proliferation genes in β-cells could be linked to the neonatal testosterone surge (Clarkson and Herbison, 2016; Corbier et al., 1992) or alternatively, mediated by locally produced testosterone in islets (Ogishima et al., 2008).

In summary, our integrative GRN analysis provides a detailed understanding of the gene regulatory mechanisms defining SC-islet and primary islet cell types. These GRNs will be a valuable resource to inform strategies for producing precision cell therapy products.

## Acknowledgements

We acknowledge support of the UCSD IGM Genomic Center and P30 DK064391 for sequencing. This work was supported by grants from the National Institutes of Health U01 DK120429 (M.S. and K.J.G.), U01 DK105541 (M.S. and K.J.G.), R01 DK068471 (M.S.), UH3 DK122639 (M.S.), and postdoctoral fellowships from the California Institute for Regenerative Medicine (H.Z.), the Diabetes Research Connection (H.Z.), and Juvenile Diabetes Research Foundation (K.V.N.). Work at the Center for Epigenomics was supported in part by the UC San Diego School of Medicine. We thank Ryan Geusz, Medhavi Mallick, and members of the Sander lab for scientific discussions and input on the project.

## Author Contributions

M.S. conceived and supervised the research; M.S., H.Z., and G.W. wrote the manuscript; H.Z., and G.W. devised experimental and computation methodologies; H.Z., K.V.N., D.K., M.M., G.G., J.K. and D.S. performed experiments; H.Z. and G.W. performed analyses of single-cell and ChIP-seq data; R.M. and K.J.G. contributed to the collection and analysis of adult human pancreas snATAC-seq and snRNA-seq data; G.G. and F.M.S. contributed immunostaining of human fetal pancreas. A.H. prepared CDX2 ChIP-seq library. D.S. and A.C.P. contributed to immunostaining of neonatal human pancreas and provided childhood human pancreas. S.P. supervised 10x single-cell assays; M.M. performed 10x single-cell assays.

## Conflict of Interest

K.J.G. does consulting for Genentech and holds stock in Vertex Pharmaceuticals.

## Methods

### RESOURCE AVAILABILITY

#### Material availability

*CDX2* knockout H1 hESC line is available upon request.

#### Data availability

Single-nucleus ATAC sequencing (snATAC-seq), Single-nucleus RNA sequencing (snATAC-seq), Single-cell RNA sequencing (scRNA-seq), and CDX2 ChIP sequencing raw and processed data are available through the Gene Expression Omnibus. Other published datasets used in this study are summarized in Supplemental Table 1a. UCSC genome browser sessions of aggregated snATAC-seq, snRNA-seq, and scRNA-seq data are available at: https://genome.ucsc.edu/s/gaowei/hg19_islet.

### EXPERIMENTAL MODEL AND SUBJECT DETAILS

#### Human pancreata and pancreatic islets

Single-cell genomic assays were performed on snap frozen pancreas tissue or isolated islets obtained from 18 adult (20 to 61 years old) and 7 childhood (13-months to 9 years old) non-diabetic donors (HbA1c ≤ 5.6) through multiple sources including: Network for Pancreatic Organ Donors with Diabetes (nPOD), Integrated Islet Distribution Program (IIDP) and Alberta Diabetes Institute (ADI) IsletCore (see **Table S1**). Islet preparations were further enriched using zinc-dithizone staining followed by hand picking, and either directly processed for single-cell RNA sequencing (scRNA-seq) or snap frozen with liquid nitrogen or dry ice. Cryosections of fixed neonatal human pancreas were obtained from nPOD. Fixed human fetal pancreatic tissue samples were provided by the MRC/Wellcome Trust-funded Human Developmental Biology Resource (HDBR; https://www.hdbr.org; stages CS20, 10 and 20 wpc; gender not established) and by the University of Washington Birth Defects Research Laboratory (stages 13, 18, 19 wpc, gender not established). Adult human pancreas tissue for immunostaining was obtained from Prodo Labs. All human tissues were obtained from de-identified donors, and protocols used in this study were approved by Institutional Review Board (IRB, protocol 091602XX) of the University of California San Diego or by the HDBR Steering Committee to the Spagnoli laboratory at King’s College London, UK (License #200523). The HDBR is a Research Ethics Committee (REC) approved and HTA licensed tissue bank.

#### Human cell culture experiments

hESC research was approved by the University of California, San Diego (UCSD), Institutional Review Board and Embryonic Stem Cell Research Oversight Committee (protocol 090165ZX).

### METHOD DETAILS

#### Maintenance and differentiation of H1 hESCs

H1 hESCs (male) were maintained as described (Geusz et al., 2021). In brief, hESCs were seeded onto Matrigel (Corning, 356238) coated tissue culture surfaces in mTeSR1 media (Stem Cell Technologies, 85850) supplemented with 1% Penicillin-Streptomycin (Thermo Fisher Scientific, 15140122), and propagated every 3 to 4 days. Accutase (Thermo Fisher Scientific, 00-4555-56) based enzymatic dissociation method was employed for passaging and 10 µM Y-27632 (Stem Cell Technologies, 72307) was supplied on the first day of each passage.

H1 hESCs were differentiated into SC-islets with a protocol we modified from previous publications (Hogrebe et al., 2020; Rezania et al., 2014; Velazco-Cruz et al., 2019). After dissociation using Accutase, H1 cells were suspended with mTeSR1 media with %1 Penicillin-Streptomycin and 10 µM Y-27632 and plated using either a 3D culture or a 2D culture condition. For the 3D culture, cells were aggregated in 5.5mL medium at a concentration of 5.5 x 10^6^ cells/well in a low attachment 6-well plate on an orbital shaker (100 rpm, 0.2 x g) in a 37 °C incubator. The following day (day 0), undifferentiated cells were washed in Stage 1/2 base medium (see below) and then differentiated using a seven-step protocol with stage-specific medium. Medium was refreshed daily until day 32. At day 8, the speed of the orbital shaker was increased to 110 rpm (0.3 x g). On day 21, cells were dissociated with Accutase, suspended in Stage 7 medium (see below) supplemented with 10 µM Y-27632 and re-aggregated at a concentration of 3 x 10^6^ cells/well in a low attachment 6-well plate on an orbital shaker (100 rpm, 0.2 x g) in a 37 °C incubator. The speed of the shaker was increased to 110 rpm (0.3 x g) on the following day.

For a subset of experiments, a 2D differentiation protocol was used which is identical to the 3D protocol with the following exceptions: H1 hESCs cells were plated onto Matrigel coated tissue culture surfaces in base medium at a concentration of 5.7 x 10^5^ cells/cm^2^. Stage 1 was extended to a total of 4 days (day 0-3). On day 29, cells were dissociated with Accutase, suspended in Stage 7 medium (see below) supplemented with 10 µM Y-27632, and re-aggregated at a concentration of 3 x 10^6^ cells/well in a low attachment 6-well plate on an orbital shaker (100 rpm, 0.2 x g) in a 37 °C incubator. The speed of the shaker was increased to 110 rpm (0.3 x g) on the following day.

Base medium for all stage-specific media was comprised of MCDB 131 medium (Thermo Fisher Scientific, 10372019) supplemented with NaHCO3 (Sigma, S6297), GlutaMAX (Thermo Fisher Scientific, 35050061), D-Glucose (Sigma, G8769), and BSA (Lampire Biological Laboratories, 7500804) using the following concentrations:

Stage 1/2 base medium: MCDB 131 medium, 1.5 g/L NaHCO3, 1X GlutaMAX, 10 mM D-Glucose, 0.5% BSA
Stage 3/4 base medium: MCDB 131 medium, 2.5 g/L NaHCO3, 1X GlutaMAX, 10 mM D-glucose, 2% BSA
Stage 5/6 base medium: MCDB 131 medium, 1.5 g/L NaHCO3, 1X GlutaMAX, 20 mM D-glucose, 2% BSA
Stage 7 base medium: MCDB 131 medium, 1.5 g/L NaHCO3, 1X GlutaMAX, 2% BSA

Media compositions for each stage were as follows:

Stage 1 (days 0−2 for 3D culture and days 0-3 for 2D culture): base medium, 100 ng/mL Activin A (R&D Systems, 338-AC/CF), 25 ng/mL Wnt3a (R&D Systems, 5036-WN, only on day 0).
Stage 2 (days 3−5 for 3D culture and days 4-6 for 2D culture): base medium, 0.25 mM L-Ascorbic Acid (Sigma, A4544), 50 ng/mL FGF7 (R&D Systems, 251-KG)
Stage 3 (days 6−7 for 3D culture and days 7-8 for 2D culture): base medium, 0.25 mM L-Ascorbic Acid, 50 ng/mL FGF7, 0.25 µM SANT-1 (Sigma, S4572), 1 µM Retinoic Acid (Sigma, R2625), 100 nM LDN193189 (Stemgent, 04-0074), 1:200 ITS-X (Thermo Fisher Scientific, 51500056), 200 nM TPB (Calbiochem, 565740)
Stage 4 (days 8−10 for 3D culture and days 9-11 for 2D culture): base medium, 0.25 mM L-Ascorbic Acid, 2 ng/mL FGF7, 0.25 µM SANT-1, 0.1 µM Retinoic Acid, 200 nM LDN193189, 1:200 ITS-X, 100 nM TPB
Stage 5 (days 11−13 for 3D culture and days 12-14 for 2D culture): base medium, 0.25 µM SANT-1, 0.05 µM RA, 100 nM LDN-193189, 1 µM T3 (Sigma, T6397), 10 µM ALK5i II (Cayman Chemicals, 14794), 10 µM ZnSO4 (Sigma, Z0251), 10 µg/mL heparin (Sigma, H3149), 1:200 ITS-X
Stage 6 (days 14−20 for 3D culture and days 15-21 for 2D culture): base medium, 100nM LDN193189, 1 µM T3, 10 μM ALK5i II, 10 μM zinc sulfate, 100 nM gamma secretase inhibitor XX (Calbiochem, 565789), 10 μg/ml heparin, 1:200 ITS-X
Stage 7 (from day 21 for 3D culture and from day 22 for 2D culture): base medium, 10 μM zinc sulfate, 10 μg/ml heparin, 1:1000 Trace Element A (Corning, 89408-312), 1:1000 Trace Element B (Corning, 89422-908), 1:100 MEM Non-Essential Amino Acids (Thermo Fisher Scientific, 11140076)

#### Dihydrotestosterone treatment

H1 hESCs were differentiated as described above. 10nM dihydrotestosterone (DHT, Sigma, D-073) was added daily to the differentiation medium starting from either day 14 (end of Stage 5) or day 21 (end of Stage 6). Methanol was used as vehicle control.

#### Generation of *CDX2* KO H1 hESC line

To generate a homozygous *CDX2* deletion H1 hESC line, sgRNAs targeting the first exons of *CDX2* were cloned into PX458 (Addgene, 48138). The plasmid was transfected into H1 hESCs with XtremeGene 9 (Roche, 6365787001), and 24 h later 5000 GFP^+^ cells were sorted into a well of six-well plate using mTeSR1 medium supplemented with 10 µM Y-27632. Individual colonies that emerged within 7 days were subsequently transferred manually into 48-well plates for expansion, genomic DNA extraction, PCR genotyping, and Sanger sequencing. A clone with a homozygous five base pair deletion in the *CDX2* coding sequence was selected. For control clones, the PX458 plasmid was transfected into H1 hESCs, and cells were subjected to the same workflow as H1 hESCs transfected with sgRNAs.

sgRNA oligos used to generate *CDX2* KO hESCs: GAGAAGCGCAGGAAGGCGCGG PCR primers used to amply edited *CDX2* genomic locus:

Forward: TCACGGCCTCAACGGTGGCT
Reverse: GCCTTCCCAAGCACCCTCCGAA

#### Flow cytometry analysis

Cell aggregates derived from hESCs were allowed to settle in microcentrifuge tubes and washed with PBS. Cell aggregates were incubated with Accutase® at 37 °C until a single-cell suspension was obtained. Cells were washed with 1 mL ice-cold flow buffer comprised of 0.2% BSA in PBS and centrifuged at 200 x *g* for 5 min. BD Cytofix/Cytoperm™ Plus Fixation/Permeabilization Solution Kit was used to fix and stain cells for flow cytometry according to the manufacturer’s instructions. Briefly, cell pellets were resuspended in ice-cold BD Fixation/Permeabilization solution (300 µL per microcentrifuge tube). Cells were incubated for 20 min at 4 °C. Cells were washed twice with 1 mL ice-cold 1X BD Perm/Wash™ Buffer and centrifuged at 4 °C and 200 x *g* or 5 min. Cells were resuspended in 50 µL ice-cold 1X BD Perm/Wash™ Buffer containing diluted antibodies, for each staining performed. Cells were incubated at 4 °C in the dark for 1–3 h. If a secondary antibody staining was required, cells were washed twice with 1 mL ice-cold 1X BD Perm/Wash™ Buffer and centrifuged at 4 °C and 200 x *g* for 5 min. Cells were resuspended in 50 µL ice-cold 1X BD Perm/Wash™ Buffer containing diluted secondary antibodies. For EdU incorporation assays coupled with the flow analysis, cells were stained for EdU prior to primary antibody staining. Cells were washed with 1.25 mL ice-cold 1X BD Wash Buffer and centrifuged at 200 x *g* for 5 min. Cell pellets were resuspended in 300 µL ice-cold flow buffer and analyzed in a FACS LSRFortessa™ system (BD Biosciences). Antibodies used were AlexaFluor® 647-conjugated anti-NKX6-1 (1:5 dilution, BD Biosciences 563338); PE-conjugated anti-insulin (1:50 dilution, Cell Signaling 8508); rabbit anti-human SLC18A1 (1:300 dilution, Sigma HPA063797); AlexaFluor® 488-conjugated donkey anti-rabbit IgG (1:1000 dilution, Jackson Immunoresearch 711-545-152); Biotin-conjugated anti-CD26 (1:500 dilution, BioLegend 302718); and Brilliant Violet 421™-conjugated Streptavidin (1:500 dilution, BioLegend 405226). Data were processed using FlowJo software v10.

#### Nucleoside analog (EdU) incorporation assay

Proliferation in SC-β-cells was assayed using Click-iT™ EdU Alexa Fluor™ 488 Flow Cytometry Assay Kit following the manufacturer’s instructions with modifications. In brief, nucleoside analog, EdU (5-ethynyl-2’-deoxyuridine, 10 μM), was added daily to SC-islets starting from day 21 until day 32 of 3D culture. Labeled cells were dissociated, fixed, and permeabilized using the same procedures as described in “Flow cytometry analysis”, and visualized with Alexa Fluor^®^ 488 azide through “click” chemistry. To detect SC-β-cell-specific EdU incorporation, cells were stained with AlexaFluor® 647-conjugated anti-NKX6-1 and PE-conjugated anti-insulin (see “Flow cytometry analysis”), and analyzed in a FACS LSRFortessa™. Data were processed using FlowJo software v10.

#### Immunofluorescence analysis

SC-islets were washed twice with PBS and fixed with 4% paraformaldehyde (PFA) for 30 min at room temperature. Fixed samples were washed twice with PBS and dehydrated in 30% (w/v) sucrose in PBS at 4 °C overnight. The following day, samples were embedded with Tissue-Tek^®^ O.C.T. Sakura® Finetek compound (VWR) in disposable embedding molds (VWR), and frozen in a dry ice-ethanol bath. Tissue blocks were sectioned at 10 µm and sections were placed on Superfrost Plus® (Thermo Fisher) microscope slides and washed twice with PBS for 10 min. On neonatal and adult human pancreas sections, we performed antigen retrieval by boiling sections in sodium citrate buffer (10 mM sodium citrate, 0.05% Tween 20, pH 6.0) for 20 min. Sections were then permeabilized with 0.1% (v/v) Triton X-100 (Sigma-Aldrich) for 30 min, and blocked with blocking buffer, consisting of 0.1% (v/v) Triton X-100 (Sigma-Aldrich) and 1% (v/v) normal donkey serum (Jackson Immuno Research Laboratories, Cat 017-000-121) in PBS for 1h at room temperature. Primary antibody incubation was conducted in the same blocking buffer at 4 °C overnight. The following day, sections were washed three times with PBS and stained with diluted secondary antibodies and Hoechst 33342 (Invitrogen, H3570) for 1h at room temperature. Stained sections were washed five times with PBS before mounting with VECTASHIELD^®^ (Vector Laboratories, H-1300). Images were obtained with a Zeiss Axio-Observer-Z1 microscope equipped with a Zeiss ApoTome and AxioCam digital camera and quantified using HALO image analysis (Indica Lab). Fetal human pancreas sections processed in the Spagnoli laboratory at King’s College London were stained with a similar procedure but using slightly different reagents, including a citrate buffer solution (Dako) for antigen retrieval, a TSA blocking buffer [0.5% TSA blocking powder (Perkin Elmer, Cat NEL 701001KT), 10% horse serum (Invitrogen, Cat 16050130) in 0.1% Triton 1x PBS] and Dako Fluorescent Mounting Medium (Dako, Cat S3023). Images were acquired on Zeiss LSM 700 laser scanning microscope.

#### Dynamic glucose-stimulated insulin secretion (GSIS) assay

GSIS assays were carried out at 37°C using the Biorep perifusion system (Biorep v5), which allows a dynamic exchange of Krebs-Ringers-Bicarbonate-HEPES (KRBH) buffer (130 mM NaCl, 5 mM KCl, 1.2 mM CaCl2, 1.2 mM MgCl2, 1.2 mM KH2PO4, 20 mM HEPES pH 7.4, 25 mM NaHCO3, and 0.1% BSA) with high (16.8 mM) and low (2.8 mM) glucose concentrations. 30-50 hand-picked SC-islets were loaded into the perifusion chamber and equilibrated with low glucose KRBH for 1h. SC-islets were then stimulated with KRBH containing indicated concentration of glucose for indicated duration. Perifusate was collected every minute, and samples from indicated time points were analyzed using STELLUX® Chemi Human C-peptide ELISA (ALPCO). SC-islets from perifusion chambers were transferred to microcentrifuge tubes and lysed by sonication for total C-peptide content measurement.

#### RNA extraction and qRT-PCR

Approximately 500 SC-islets were collected and washed before RNA isolation using the RNeasy Micro kit (QIAGEN) according to the manufacturer’s instructions. RT-qPCR was performed as previously described (Wortham et al., 2019). In brief, 500 ng for total RNA was converted to cDNA using iScript™ cDNA Synthesis Kit (Bio-Rad). Gene expression was quantified with iQ™ SYBR® Green Supermix (Bio-Rad). Primers used for qPCR were listed below:

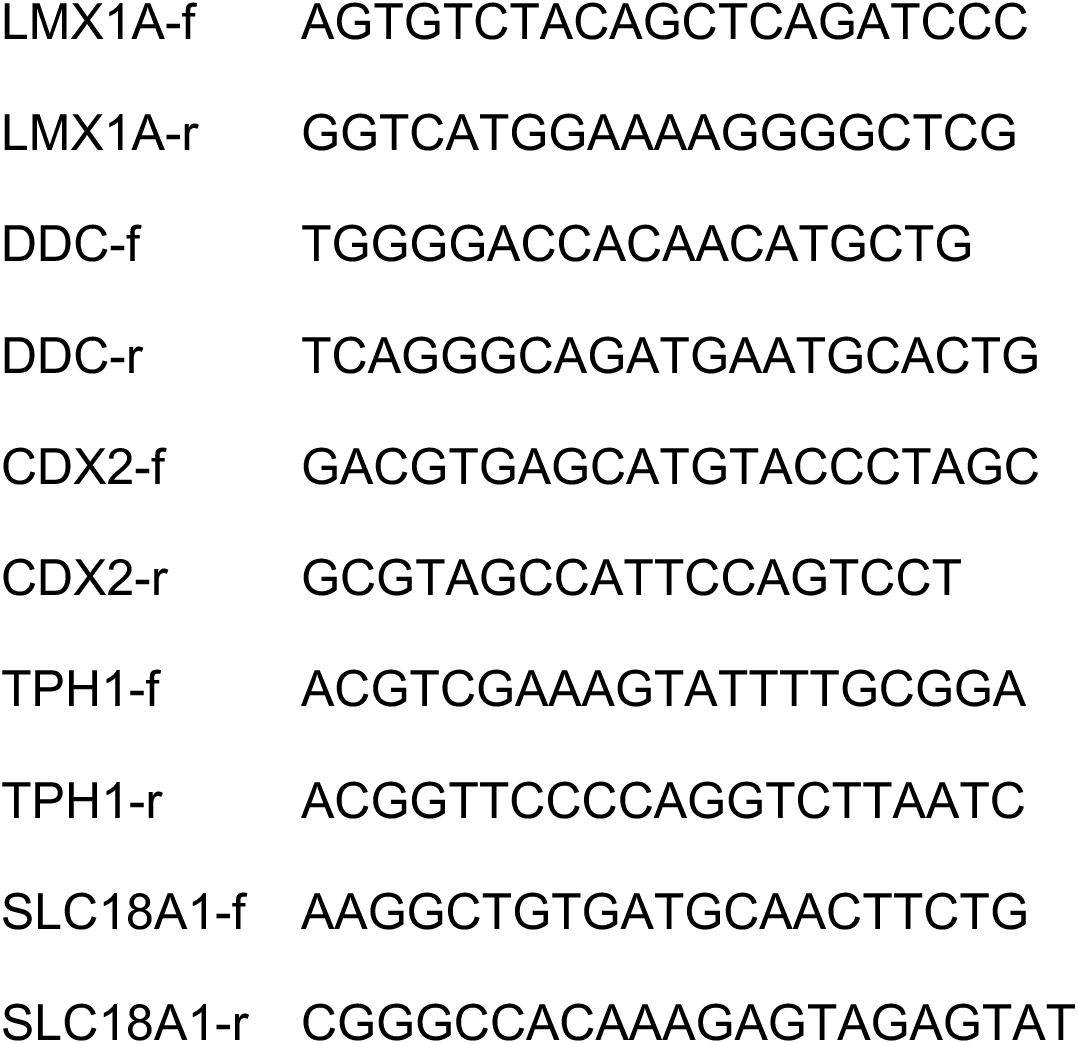

#### Single-cell RNA-sequencing (scRNA-seq)

Differentiating aggregates and hand-pick human islets were collected in microcentrifuge tubes and washed with PBS. Accutase^®^ was used to dissociate aggregates into single cells, which were then stained with propidium iodide (Sigma) in a PBS solution containing 0.2% BSA. Approximately 200,000 live cells (propidium iodide-negative) were sorted with a FACSAriaTM Fusion Flow Sorter at a sorting speed lower than 3,000 events per second to minimize damage to the cells. Sorted cells were pelleted with 250 x g for 5 minutes at 4°C, and counted with a Scepter™ automated cell counter. 10,000 accurately counted cells per sample were loaded onto a 10X Chromium Controller for GEM formation and cell barcoding using Next GEM Single Cell 3’ v3.1 reagents. Barcoded single cells were subjected to cDNA synthesis and sequencing library construction using 10X Next GEM Single Cell 3’ v3.1 reagents according to manufacturer’s instructions. Final libraries were quantified using a Qubit fluorimeter (Life Technologies) and the fragmented cDNA was verified using a Tapestation (High Sensitivity D1000, Agilent). Libraries were sequenced on NextSeq 500, HiSeq 4000 or NovaSeq 6000 sequencers (Illumina) and reads were trimmed afterwards to fit into corresponding analysis pipeline.

#### Single-nucleus RNA-sequencing (snRNA-seq)

Nuclei were isolated from approximately 1,000 differentiated aggregates (∼1,000 cells per aggregate) or approximately 35 mg of frozen human pancreas using a nuclei permeabilization buffer [0.1% Triton X-100 (Sigma-Aldrich, T8787), 1X Pierce Protease Inhibitor (Fischer, PIA32965), 1 mM DTT (Sigma-Aldrich, D9779), Recombinant RNase inhibitor (0.2 U/μl; Promega, 2% Fatty-acid-free BSA in PBS (Proliant, 7500804; Corning, 21-040-CV)]. For differentiated aggregates, nuclei extraction was done in a glass dounce, and for frozen human pancreas, samples were pulverized and resuspended in the nuclei permeabilization buffer.

Samples were incubated on a rotator for 5 min at 4°C and then centrifuged at 500g for 5 min (Eppendorf, 5920R; 4°C, ramp speed of 3/3). Supernatant was removed and pellet was resuspended in sort buffer [1mM EDTA (Invitrogen, 15575020), 0.2U/μL Recombinant RNAsin (Promega, PAN2515), 1% Fatty-acid-free BSA in PBS (Proliant, 7500804; Corning, 21-040-CV) and stained with DRAQ7 (1:150; Cell Signaling Technology, 7406). 60,000 nuclei were sorted using an SH800 sorter (Sony) into 50 μl of collection buffer [1.0U/μL Recombinant RNAsin (Promega, PAN2515), 5% Fatty-acid-free BSA in PBS (Proliant, 7500804; Corning, 21-040-CV)]. Sorted nuclei were then centrifuged at 1000 g for 15 min (Eppendorf, 5920R; 4°C, ramp speed of 3/3), and supernatant was removed. Nuclei were resuspended in reaction buffer [RNase inhibitor (0.2U/μL Recombinant RNAsin (Promega, PAN2515), 1% Fatty-acid-free BSA in PBS (Proliant, 7500804; Corning, 21-040-CV) and counted using a hemocytometer.

16,550 nuclei were loaded onto a Chromium controller (10x Genomics). Libraries were generated using the Chromium Next GEM Single Cell 3’ GEM, Library & Gel Bead Kit v3.1 (10x Genomics, PN-1000121) according to the manufacturer specifications. Complementary DNA was amplified for 12 PCR cycles. SPRISelect reagent (Beckman Coulter) was used for size selection and cleanup steps. Final library concentration was assessed by the Qubit dsDNA HS Assay Kit (Thermo Fisher Scientific), and fragment size was checked using TapeStation High Sensitivity D1000 (Agilent) to ensure that fragment sizes were distributed normally around 500 bp. Libraries were sequenced using a NextSeq 500 or NovaSeq 6000 (Illumina).

#### Single-nucleus ATAC-sequencing (snATAC-seq)

Nuclei extraction and sorting were done using the same methodology as described in “Single nucleus RNA-seq”. Single nucleus ATAC-seq libraries were generated using either the Chromium Chip E Single Cell ATAC Kit (10x Genomics, 1000086) or Chromium Next GEM Single Cell ATAC Library & Gel Bead Kit v1.0 (10x Genomics, 1000175) with Chromium Next GEM Chip H Single Cell Kit (1000161) following the manufacturer’s instructions. Indexes used were Chromium i7 Multiplex Kit N, Set A (10x Genomics, 1000084) and Single Index Kit N Set A (1000212)10x Genomics, 1000084), respectively. Final libraries were quantified using a Qubit fluorimeter (Life technologies) and the nucleosomal pattern was verified using a Tapestation (High Sensitivity D1000, Agilent). Libraries were sequenced on NextSeq 500, HiSeq 4000 or NovaSeq 6000 sequencers (Illumina) and reads were trimmed afterwards to fit into corresponding analysis pipeline.

#### Chromatin immunoprecipitation sequencing (ChIP-seq)

ChIP-seq was performed using the ChIP-IT High-Sensitivity kit (Active Motif) according to the manufacturer’s instructions. Briefly, from day 21 SC-islets 5–10 × 10^6^cells were harvested and fixed on a rocker for 15 min in an 11.1% formaldehyde solution. The reaction was quenched for 5 min in 0.125 M glycine, cells washed in DPBS containing 0.5% NP-40, then once again in DPBS supplemented with 0.5% NP-40 and 1 mM PMSF. Cells were lysed by sonication with a Bioruptor® Plus (Diagenode), on high for 3 × 5 min (30 s on, 30 s off). 30 µg of the resulting sheared chromatin was used for each immunoprecipitation. Equal quantities of sheared chromatin from each sample were used for immunoprecipitations carried out at the same time. 6 μg anti-CDX2 antibody (A300-691A, Bethyl Laboratories) was used for the ChIP-seq assay. Chromatin was incubated with primary antibody overnight at 4 °C on a rotator followed by incubation with Protein G agarose beads for 3 h at 4 °C on a rotator. Reversal of crosslinks and DNA purification were performed according to the ChIP-IT High-Sensitivity instructions, with the modification of incubation at 65 °C for 2-3 h, rather than at 80 °C for 2 h. Sequencing libraries were constructed using KAPA DNA Library Preparation Kits for Illumina® (Kapa Biosystems) and library sequencing was performed on either a HiSeq 4000 System (Illumina®) or NovaSeq 6000 System (Illumina®) with single-end reads of either 50 or 75 base pairs (bp). Sequencing was performed by the UCSD Institute for Genomic Medicine (IGM) core research facility. For the ChIP-seq experiment, replicates from two independent hESC differentiations were generated.

#### ChIP-seq data analysis

Bowtie2 (Langmead and Salzberg, 2012) (v2.3.4.1) was used for mapping of raw data to the human reference genome hg19 with a maximum of 2 mismatches allowed in the seed region, discarding reads aligning to multiple sites. Duplicate reads were removed using SAMtools (Li et al., 2009). DeepTools (Ramirez et al., 2014) was used to generate bigwig format tracks for visualization in UCSC Genome Browser. Peak calling was performed using MACS2 (Zhang et al., 2008) with default setting for TF ChIP-seq and ChIP-seq input as the background control.

#### Single-cell raw data processing and quality control

##### Data processing using Cell Ranger software

Alignment to the hg19 genome and initial processing were performed using the 10x Genomics Cell Ranger ATAC v1.1.0 and Cell Ranger RNA v.3.0.2 pipelines. Sample information and a summary of the Cell Ranger ATAC-seq and RNA-seq quality metrics are provided in **Table S1**.

##### Filtering barcode doublets and low-quality cells for each individual donor

Cell barcodes from the 10x Chromium snATAC-seq assay may have barcode multiplets that have more than one oligonucleotide sequence (Lareau et al., 2020). We used ‘clean_barcode_multiplets_1.1.py’ script from 10x to identify barcode multiplets for each donor and excluded these barcodes from further analysis. We then filtered low quality snATAC-seq profiles by total UMIs (<1,000), fraction of reads overlapping TSS (<15%), fraction of reads overlapping called peaks (<30%), and fraction of reads overlapping mitochondrial DNA (>10%) according to the distribution of these metrics for all barcodes. We also excluded profiles that had extremely high unique nuclear reads (top 1%), fraction of reads overlapping TSS (top 1%) and called peaks (top 1%) to minimize the contribution of these barcodes to our analysis. For RNA-seq data, we used total UMIs (<1,000) and fraction of reads overlapping mitochondrial DNA (>10%) to filter cells with low quality RNA profiles. We also excluded profiles that had extremely high total UMIs (top 1%) to minimize the contribution of these barcodes to our analysis.

##### Cell clustering

After filtering low quality cells, we checked ATAC and RNA data quality from each sample by performing an initial clustering using Scanpy (v.1.6.0) (Wolf et al., 2018). For ATAC-seq data, we partitioned the hg19 genome into 5 kb sliding windows and removing windows overlapping blacklisted regions from ENCODE (Amemiya et al., 2019; Consortium, 2012) (https://www.encodeproject.org/annotations/ENCSR636HFF/). Using 5 kb sliding windows as features, we produced a barcode-by-feature count matrix consisting of the counts of reads within each feature region for each barcode. Detailed pipeline to process ATAC-seq data can be found in our previous work (Chiou et al., 2021). We normalized each barcode to a uniform read depth and extracted highly variable features. Then, we regressed out the total read depth for each cell, performed PCA, and extracted the top 50 principal components to calculate the nearest 30 neighbors using the cosine metric, which were subsequently used for UMAP dimensionality reduction with the parameters ‘min_dist=0.3’ and Leiden (Traag et al., 2019) clustering with the parameters ‘resolution=0.8’.

We then performed initial cell clustering for cells from all donors using similar methods to cluster cells for each donor. Of note, we extracted highly variable features across cells from all experiments. Since read depth was a technical covariate specific to each experiment, we regressed this out on a per-experiment basis. We also used Harmony (Korsunsky et al., 2019) to adjust for batch effects across experiments. We identified clusters and subclusters (‘resolution’=1.5) with significantly different total UMIs, fraction of reads overlapping TSS, or fraction of reads overlapping called peaks compared to other clusters and subclusters. We excluded these clusters and subclusters and obtained final cell clusters by performing cell clustering using identical methods for initial clustering of all cells. We determined the cell type represented by each cluster by examining chromatin accessibility at the promoter regions of known marker genes.

#### Generating fixed-width and non-overlapping peaks that represent cCREs across cell types

We called peaks for each cell type using the MACS2 call peak command with parameters ‘--nomodel --extsize 200 –shift 0 --keep-dup all -q 0.05’ and filtered these peaks by the ENCODE hg19 blacklist. For each cell type, we generated fixed-width peaks (summits of these peaks from macs2 were extended by 250 bp on either side to a final width of 501 bp), as previously described (Satpathy et al., 2019). We quantified the significance of these fixed-width peaks in each cell type by converting the MACS2 peak scores (−log10(Q value)) to a ‘score quantile’. Then, fixed-width peaks for each cell type were combined into a cumulative peak set. As there are overlapping peaks across cell types, we retained the most significant peak and any peak that directly overlapped with that significant peak was removed. This process was iterated to the next most significant peak and so on until all peaks were either kept or removed due to direct overlap with a more significant peak. These fixed-width and non-overlapping peaks were defined as candidate *cis*-regulatory element, or cCREs.

#### Differential gene expression analysis

We used generalized linear regression model (glm function in R) to call differential expressed genes between different cell types or states from snRNA-seq data. Expression level of genes were normalized by total count for individual cells. In addition to cell types or states annotation, we also considered total count of individual cells as covariate in the model to calculate coefficient and p value. Adjusted p values (FDRs) were obtained using p.adjust function in R with a Benjamini & Hochberg method. Differentially expressed genes were selected using FDR<0.05 as cutoff.

#### K-means clustering

For α-cells, we used the cell type-by-genes count matrix and differentially expressed genes between α-cells from SC-islets, childhood, and adult primary islets (FDR<0.05) as input. We normalized the expression level of genes using total counts and performed K-means clustering analysis using kmeans function in R. We then repeated the same procedure for β-cells.

#### Integrating snATAC-seq and sc/snRNA-seq data

We used the Seurat package (Stuart et al., 2019) to integrate single modality snATAC-seq and sc/sn RNA-seq datasets (https://satijalab.org/seurat/articles/atacseq_integration_vignette.html). Raw count matrices of snATAC-seq and sc/snRNA-seq data, as well as cell clustering results were loaded into the Seurat package as input. To integrate and establish connections between transcriptome (sc/sn RNA-seq) and accessible chromatin (snATAC-seq) profiles, we first inferred gene activity scores from snATAC-seq data using the GeneActivity function and performed log-normalization. Gene activity scores from snATAC-seq data and gene expression from sc/snRNA-seq data were then compared and linked using FindTransferAnchors function with a Canonical Correlation Analysis (CCA) based dimension reduction method. Using anchors identified in the CCA space, cluster identities and mRNA counts of the snRNA-seq dataset were transferred to cells in snATAC-seq datasets. We applied the procedure to integrate snATAC- and sc/snRNA-seq data of SC-islets and endocrine cells from SC-islets and primary endocrine cells from human pancreas.

#### Integrating SC-islet endocrine cells with primary human pancreatic endocrine cells

We performed separate integration analyses for snATAC-seq data and sc/snRNA-seq data obtained from SC-islet endocrine cells and primary human pancreatic endocrine cells. For snATAC-seq, processed matrices from Scanpy were imported into the “Signac” R package (https://satijalab.org/signac/articles/pbmc_vignette.html). Data from SC- and primary endocrine cells were merged and normalized using a frequency-inverse document frequency (TF-IDF) method and dimension reduction was performed using singular value decomposition (SVD) followed by latent semantic indexing (LSI). Cells from the two datasets were compared in the LSI space and integration anchors were identified using the FindIntegrationAnchors function, and integrated with those integration anchors using IntegrateData. For sc/snRNA-seq datasets, integration was performed following “Seurat” data integration instructions (https://satijalb.org/seurat/articles/integration_introduction.html). In brief, SC- and primary endocrine cells were imported into “Seurat” package from “Scanpy” with original dimension reductions (PCA and UMAP) remaining the same. Integration anchors were found by comparing datasets in the PCA space using FindIntegrationAnchors before datasets were integrated using IntegrateData. Corrected snATAC-seq and sc/snRNA-seq matrices after integration were normalized and dimensionally reduced. Cells from SC- and primary endocrine cells were co-embedded on a same UMAP following the method described in “Single-cell raw data processing and quality control”. Both original cell identities and new identities obtained after integration were visualized using the first two UMAP components and compared.

#### TF motif enrichment analysis

Using the barcode-by-peaks (501 bp fixed-width peaks) count matrix as input, we inferred enrichment of TF motifs for each barcode using chromVAR (Schep et al., 2017) (v.1.4.1). We filtered cells with minimal reads less than 1500 (min_depth=1500) and peaks with fraction of reads less than 0.15 (min_in_peaks=0.15) by using ‘filterSamplesPlot’ function from chromVAR. We also corrected GC bias based on ‘BSgenome.Hsapiens.UCSC.hg19’ using the ‘addGCBias’ function. Then, we used the TF binding profiles database JASPAR 2020 motifs (Fornes et al., 2020) and calculated the deviation z-scores for each TF motif in each cell by using the ‘computeDeviations’ function. High-variance TF motifs across all cell types were selected using the ‘computeVariability’ function with the cut-off 1.15 (*n*=315). For each of these variable motifs, we calculated the mean z-score for each cell type and normalized the values to 0 (minimal) and 1 (maximal).

#### Building pseudotime trajectories with Monocle3

For each developmental trajectory, cells from indicated lineages were selected in the snATAC-seq dataset Seurat object using the subset function and the subset object was imported into Monocle3 (Qiu et al., 2017) using the as.cell_data_set functions with default settings. snATAC-seq UMAP coordinates were used to estimate distance between cells and to identify the nearest neighbor cell. This process was combined with the establishment of a lineage trajectory using the learn_graph function with close_loop = F, and learn_graph_control=list(ncenter=500,minimal_branch_len=10). Pseudotime trajectory roots were chosen empirically based on prior knowledge of pancreas development, with an interactive interface using the order_cells function. To minimize computational noise introduced by the sparse nature of single-cell data, we created pseudo-bulk samples by cutting the entire pseudotime trajectory into 12 pseudotime bins and aggregated cells within each bin using the aggregate_by_cell_bin function. We then integrated chromatin accessibility, gene expression and TF motif enrichment data into each single-cell to compute CPM values of cCREs, genes, and mean motif enrichment scores in each pseudotime bin. Values were then scaled and plotted on a heatmap for visualization.

#### Inferring gene regulatory networks (GRNs)

##### Computing correlation between cCRE accessibility and target gene expression

To identify putative target genes of cCREs, we combined and modified previously published methods (Li et al., 2021). First, we identified cCRE-gene pairs with physical interaction with the following three methods: 1. cCREs within ± 1 kb of a TSS were defined as gene promoter cCREs. Promoter-gene pairs were established across all expressed genes in each cell type. 2. cCREs located outside ± 1 kb, but within ± 50 kb of a TSS were classified as proximal elements, and all proximal cCRE-gene interactions in each cell type were considered. 3. We used Cicero (Pliner et al., 2018) to calculate co-accessibility between long distance cCREs (see “Computing co-accessibility using Cicero”) and identified distal cCRE-gene pairs for individual cell types.

We then generated pseudo-bulk ATAC and mRNA profiles by aggregating single cells of the same cell type from different cell sources (stem cell-derived, endocrine cells from childhood or adult pancreas) and collection times during SC-islet differentiation (D11, D14, D21, D32, D39). In total there were 16 pseudo-bulk ATAC and RNA profiles. CPM (counts per million reads) values of cCRE accessibility and gene expression in each pseudo-bulk ATAC and RNA profile were calculated, cCREs with low accessibility (maximum CPM value across pseudo-bulk ATAC profile <1) and gene with low expression genes (maximum CPM value across pseudo-bulk RNA profile <3) were excluded from further analysis. Finally, we calculated the Spearman correlation coefficient (SCC) between cCRE accessibility and target gene expression across all cCRE-gene pairs identified above. To estimate background, we generated permuted pseudo-bulk ATAC and RNA profiles by randomly shuffling identities of pseudo-bulk profiles, cCREs, and genes. We estimated False-positive detection rates (FDR) (Zhu et al., 2021) based on the fraction of detected pairs from the shuffled group. Empirically defined cutoffs were used to identify the final lists of cCRE-gene pairs.

##### Computing co-accessibility using Cicero

We used Cicero (Pliner et al., 2018) (v.1.3.4.10) to calculate co-accessibility scores for pairs of peaks in each individual cell type. Using SC-β-cell as example, we started from the merged peak by cell sparse binary matrix, extracted SC-β-cells, and filtered out peaks that were not present in SC-β-cells. We used the ‘make_cicero_cds’ function to aggregate cells based on the 50 nearest neighbors. We then used Cicero to calculate co-accessibility scores using a window size of 1 Mb and a distance constraint of 250 kb. We then repeated the same procedure for other cell types. We used a co-accessibility threshold of 0.05 to define pairs of peaks as co-accessible. Peaks within and outside ± 5 kb of a TSS in GENCODE V19 were considered proximal and distal, respectively. Peaks within ± 500 bp of a TSS in GENCODE V19 were defined as promoter. Co-accessible pairs were assigned to one of three groups: distal-to-distal, distal-to-proximal and proximal-to-proximal. Distal-to-proximal co-accessible pairs were defined as potential enhancer-promoter connections. Genes linked to proximal or distal cCREs were identified.

##### Computing correlation between transcription factor (TF) expression and cCRE accessibility

We used a position frequency matrix (PFMatrixList object) of TF DNA-binding preferences from the JASPAR 2020 database (Fornes et al., 2020) and width-fixed peaks as input to perform TF footprinting analysis. We used the ‘matchMotifs’ function in the R package motifmatchr to infer cCREs bound by TFs. This analysis established a preliminary set of TF-cCRE pairs. A matching set of aggregated pseudo-bulk ATAC and RNA profiles (see “Computing correlation between cCRE accessibility and target gene expression”) was used to quantify CPM values of TF expression and cCRE accessibility. We then calculated SCCs across all pseudo-bulk and permuted pseudo-bulk aggregates through randomization. The FDR was calculated using the same method as described in “Computing correlation between cCRE accessibility and target gene expression” and empirically defined cutoffs were used to define significantly correlated TF-cCRE pairs.

##### Establishment of cell type-specific GRNs

We identified highly variable cCREs across cell types based on cCRE-by-pseudo-bulk count matrices. We then performed k-means clustering of highly variable cCREs to identify cCREs modules, defined by cCREs exhibiting a similar accessibility pattern across cell types. Cell type-specific cCRE modules and cell type-shared cCRE modules were identified. Upstream TFs and downstream target genes of each cCRE from each module were used to define cell type-specific GRNs. To visualize features of cell type-specific GRNs, we performed a UMAP (umap function in “uwot” package in R) based dimension reduction analysis of the pseudocell-by-cCRE accessibility matrices used for the correlation-based GRN inference, and plotted individual cCREs using the first two UMAP components. Onto this cCRE UMAP, we plotted different features of cCREs used in the GRN analysis and retained cCRE module-specific information in the plots. Those cCRE features include: 1. Accessibility in each pseudocell; 2. cCRE module identity; 3. Correlation (SCC) between each cCRE and a given TF; 4. cCREs co-bound by two TFs; 5. cCRE pseudotime values.

##### Identification of cell type-specific transcriptional regulators

We used Fisher’s exact test to identify cell type-specific TFs. For each TF in query, we computed (fisher.test function in R) odds ratio and p value describing enrichment of cCREs bound by the TF within a cCRE module compared to TF-bound cCREs across all cCRE modules. This process was repeated for all TFs in all cCRE modules and adjusted p values (FDR) were obtained using p.adjust function in R with a Benjamini & Hochberg method. Significantly enriched TFs were selected if FDR<0.05.

##### Inferring cell type-specific TF interactions

TF interactions were inferred by one TF (TF1) binding to the same set of cCREs also bound by another TF (TF2) in a cell type-specific cCRE module. To test this, we focused on one cCRE module each time and estimated the enrichment of TF1-bound cCREs within TF2 binding sites compared to those in the entire cCRE module. This enrichment was summarized using odds ratios and p values calculated with fisher.test in R. This procedure was repeated for all TF pairs in all cCRE modules and adjusted p values (FDR) were obtained using p.adjust function in R with a Benjamini & Hochberg method. Significantly TF interactions were selected if FDR<0.05.

##### Pseudotime ordering of transcriptional programs

Transcriptional programs were ordered separately for α-cell, β-cell and SC-EC lineages. In each lineage, cells were ordered on a pseudotime trajectory established in Monocle3 (described in “Building pseudotime trajectories with Monocle3”). Both cCRE accessibility and gene expression levels were plotted in each single cell on the pseudotime trajectory. Since chromatin accessibility signals are binary, for each cCRE, we estimated the density of accessible cCREs along pseudotime using the density function in R, and identified pseudotime points with highest cCRE density. For each gene, we fitted gene expression along pseudotime with the smooth.spline function, and identified pseudotime points with maximum gene expression. cCRE accessibility and gene expression pseudotime values were defined as time points with highest density of accessible cCREs and maximum gene expression, respectively. Based on these pseudotime values, cCREs and genes were aligned and ordered for each lineage. Using the established GRN that connects TFs to cCREs and target genes, we were able to plot entire transcriptional programs downstream of each TF in a lineage-specific manner.

#### Gene ontology (GO) enrichment analysis

We performed gene ontology and pathway enrichment analysis using R package Enrichr (Kuleshov et al., 2016). Libraries “GO_Biological_Process_2018”, “GO_Cellular_Component_2018”, “GO_Biological_Process_2018”, “KEGG_2019_Human”, “MSigDB_Hallmark_2020”, “Reactome_2016” were used with default parameters. To compare enrichment among multiple gene sets, GO and pathway terms significantly enriched (p value<0.05) in at least one gene set were merged. Odds ratios and p values of those terms in each gene set were summarized in a dot plot.

### QUANTIFICATION AND STATISTICAL ANALYSIS

Statistical analyses were performed using GraphPad Prism (v8.1.2), and R (v3.6.1). Statistical parameters such as the value of n, mean, standard deviation (SD), p values, and the statistical tests used are reported in the figures and figure legends. In H1 hESC differentiation experiments, the “n” refers to the number of independent hESC differentiation experiments analyzed (biological replicates). In human pancreas immunofluorescence staining “n” indicates the number of donors from which samples were obtained. All bar graphs and line graphs are displayed as mean ± SD. Paired (if observations were related) or unpaired (observations were independent) student’s t-tests were used for two-sample comparisons. For multiple-sample comparisons and comparisons done between multiple types of variables, one-way and two-way ANOVA was used, respectively. ANOVAs were coupled with one of the three multiple-comparisons tests: Šidák-Holm’s test (for two column data in two-way ANOVA); Tukey (if every column compared with every other column); or Dunnett’s test (if every column compared with a control column).

## Supplemental Figures

**Supplemental Figure 1.**
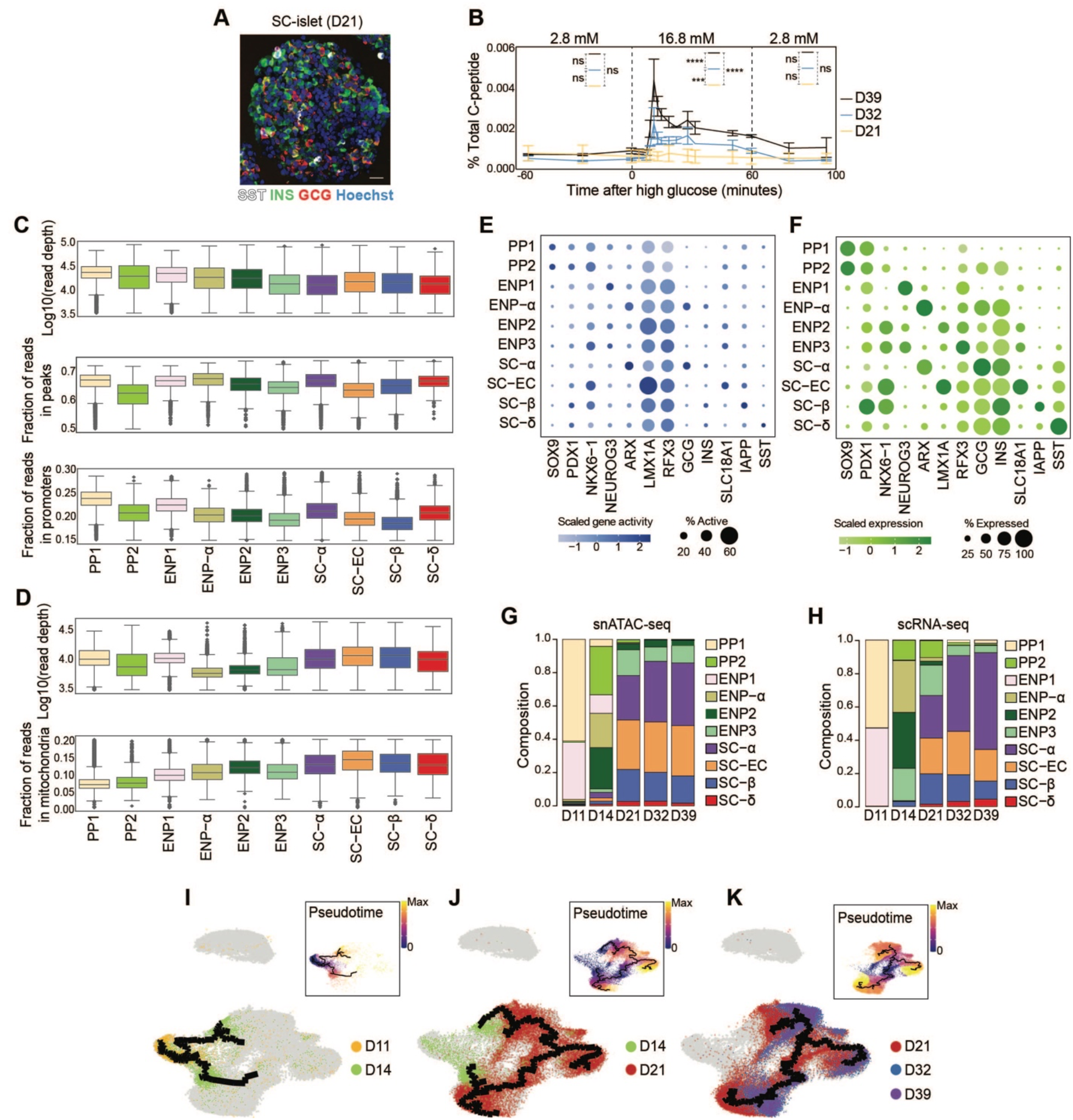
Quality control of snATAC-seq and scRNA-seq data analysis. (A) Representative immunofluorescent images for somatostatin (SST), insulin (INS) and glucagon (GCG) at D21 of SC-islet differentiation. Nuclei were labeled with Hoechst. Scale bar, 20 μm. (B) Human C-peptide secretion by SC-islets at D21, D32 and D39 during perifusion with the indicated glucose (Glc) concentrations (in mM). Data are shown as mean ± S.D. at the indicated time points. ns, not significant, \*\*\**P* < 0.001, \*\*\*\**P* < 0.0001, using two-way ANOVA followed by Tukey’s multiple comparisons test for different days of differentiation in each time block. (C, D) Box plots of quality control matrices for snATAC-seq (C) and scRNA-seq (D) in each cluster in Figure 1b, showing that these metrics do not drive single-cell grouping in UMAP space. (E, F) Dot plots showing scaled average gene activity (E) or gene expression (F). The color of each dot represents the average gene activity (E) or expression level (F) and the size of each dot the percentage of positive cells for each gene. (G, H) Relative abundance of each cell type based on snATAC-seq (G) and scRNA-seq (H) UMAP annotations in Figure 1B. Columns represent SC-islet differentiation time points. (I-K) Trajectory analysis based on chromatin accessibility, showing trajectories from D11 and D14 (I), D14 and D21 (J), and D21 and D32/39 (K) data with ENP1, ENP2, and ENP3 set as the root, respectively. Cells were color-coded by either time point of collection or pseudotime values (insets). PP1 and PP2 cells were excluded from the analysis.

**Supplemental Figure 2.**
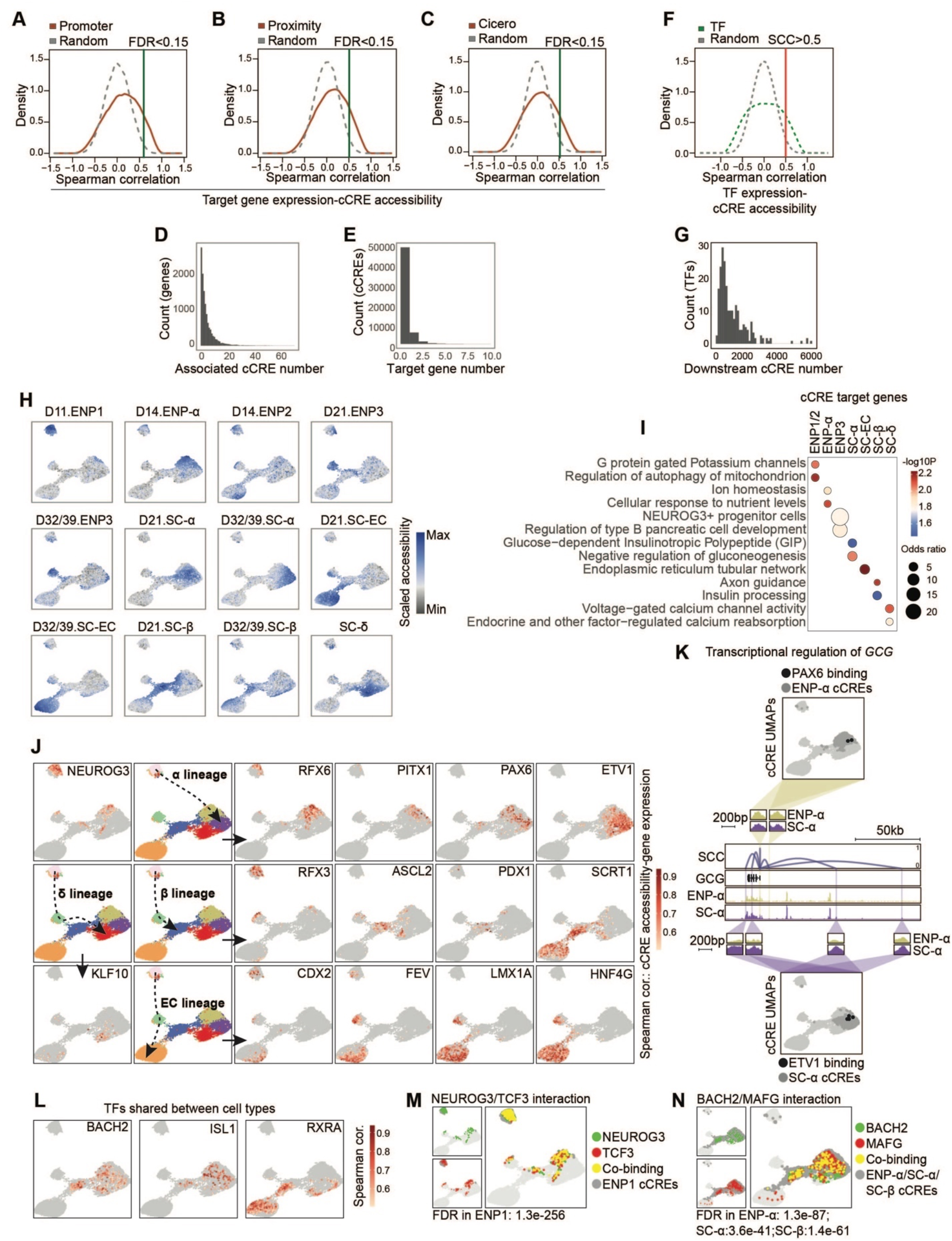
Building a gene regulatory network of stem cell islet development. (A-C) Identification of putative cCRE-gene pairs. cCREs were assigned to a gene if they are located (A) in promoter (± 500 bp) of target gene; (B) in proximity (± 50 kb) of the gene promoter; or (C) show co-accessibility with the gene promoter by Cicero analysis. A total of 65,479 positively correlated cCRE-gene pairs were identified using an empirically defined significance threshold of FDR<0.15. (D, E) Histogram showing characteristics of cCRE-gene pairs. Each target gene in the GRN is regulated by a mean of 5.3 cCREs (D) and each cCRE regulates a mean of 1.2 target genes (E). (F) Identification of putative TF-cCRE pairs. Putative TF-cCRE pairs were defined using motifmatchr. A total of 280,019 significantly correlated TF-cCRE pairs were identified using an empirically defined threshold of a spearman correlation coefficient (SCC)>0.5. (G) Histogram showing characteristics of TF-cCRE pairs. Each TF has a mean of 1,053 predicted binding sites. (H) UMAP projections of scaled chromatin accessibility of all cCREs in each aggregated pseudo-cell. Pseudo-cells were aggregated based on time point of differentiation and assigned cell type identity. (I) Enriched gene ontology terms/pathways among target genes associated with each cCRE module. Significance (-log10 p-value) and odds ratio of the enrichments are represented by color and dot size, respectively. (J) UMAP projections of TF-cCRE correlations for select cell type-enriched TFs. Spearman correlation coefficients between TF expression and cCRE accessibility are shown. (K) UMAP locations and genome browser snapshots of predicted PAX6- or ETV1-bound cCREs at the *GCG* gene locus. Dark grey dots indicate cCRE modules with predicted TF interactions. Genome browser tracks show aggregated ATAC reads in ENP-α and SC-α-cells. All tracks are scaled to uniform 1×10^6^ read depth. SCC, spearman correlation coefficients for cCRE accessibility and target gene expression. (L) UMAP projections of TF-cCRE correlations for select TFs shared between different cCRE module. Spearman correlation coefficients between TF expression and cCRE accessibility are shown. (M, N) UMAP projections of predicted TF-TF interactions. Green dots, cCREs bound by background TF; red dots, cCREs bound by test TF; yellow dots, cCREs co-bound by both TFs; dark grey dots, cCRE module(s) with predicted TF interaction(s).

**Supplemental Figure 3.**
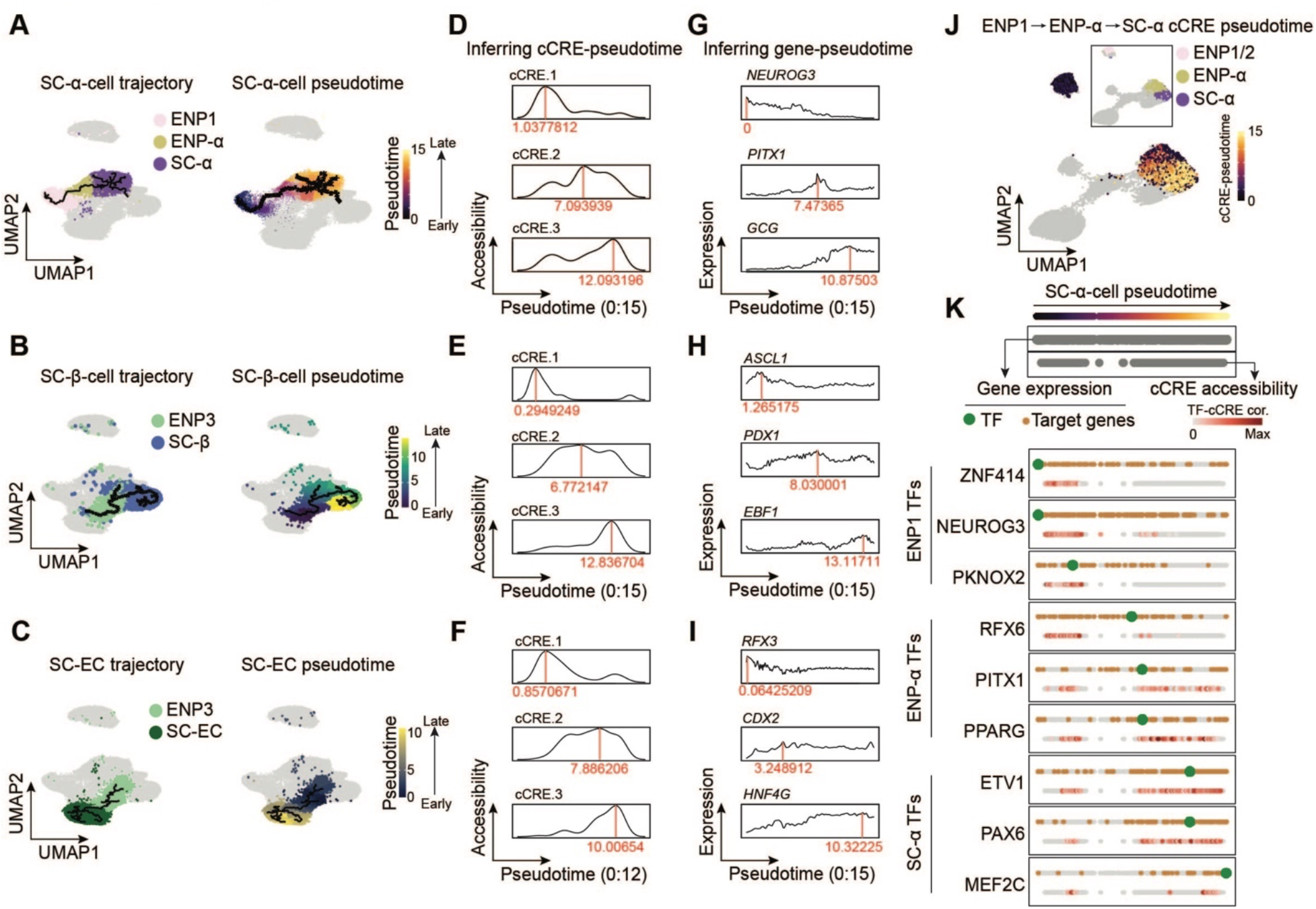
Pseudotime ordering of lineage-specific transcriptional programs. (A-C) Feature plots showing cell cluster identity (left) and pseudotime values (right) on snATAC-seq UMAPs. (A) SC-α-cell trajectory (ENP1/ENP-α/SC-α-cells), (B) SC-β-cell trajectory (ENP3/SC-β-cells) and (C) SC-EC trajectory (ENP3/SC-ECs) are shown. (D-F) Representative illustration of inference of cCRE accessibility in pseudotime on SC-α-cell (D), SC-β-cell (E) and SC-EC (F) trajectories. Values in red represent cCRE pseudotime values based on maximal cCRE accessibility along the trajectory. (G-I) Representative illustration of inference of gene expression in pseudotime on SC-α-cell (G), SC-β-cell (H) and SC-EC (I) trajectories. Values in red represent gene pseudotime values based on maximal gene expression along the trajectory. (J) UMAP projections of cCRE pseudotime on SC-α-cell lineage trajectory. Insets show cell type annotations of cCRE modules. (K) Pseudotime ordering of transcriptional programs along SC-α-cell lineage trajectory from ENP1 and ENP-α progenitors. Gene expression and cCRE accessibility were assigned pseudotime values and plotted in two separate dotted lines (genes, top; cCREs, bottom). For each shown TF, the TF (green), TF-bound cCREs (colored based TF-cCRE correlations) and target genes (brown) are shown.

**Supplemental Figure 4.**
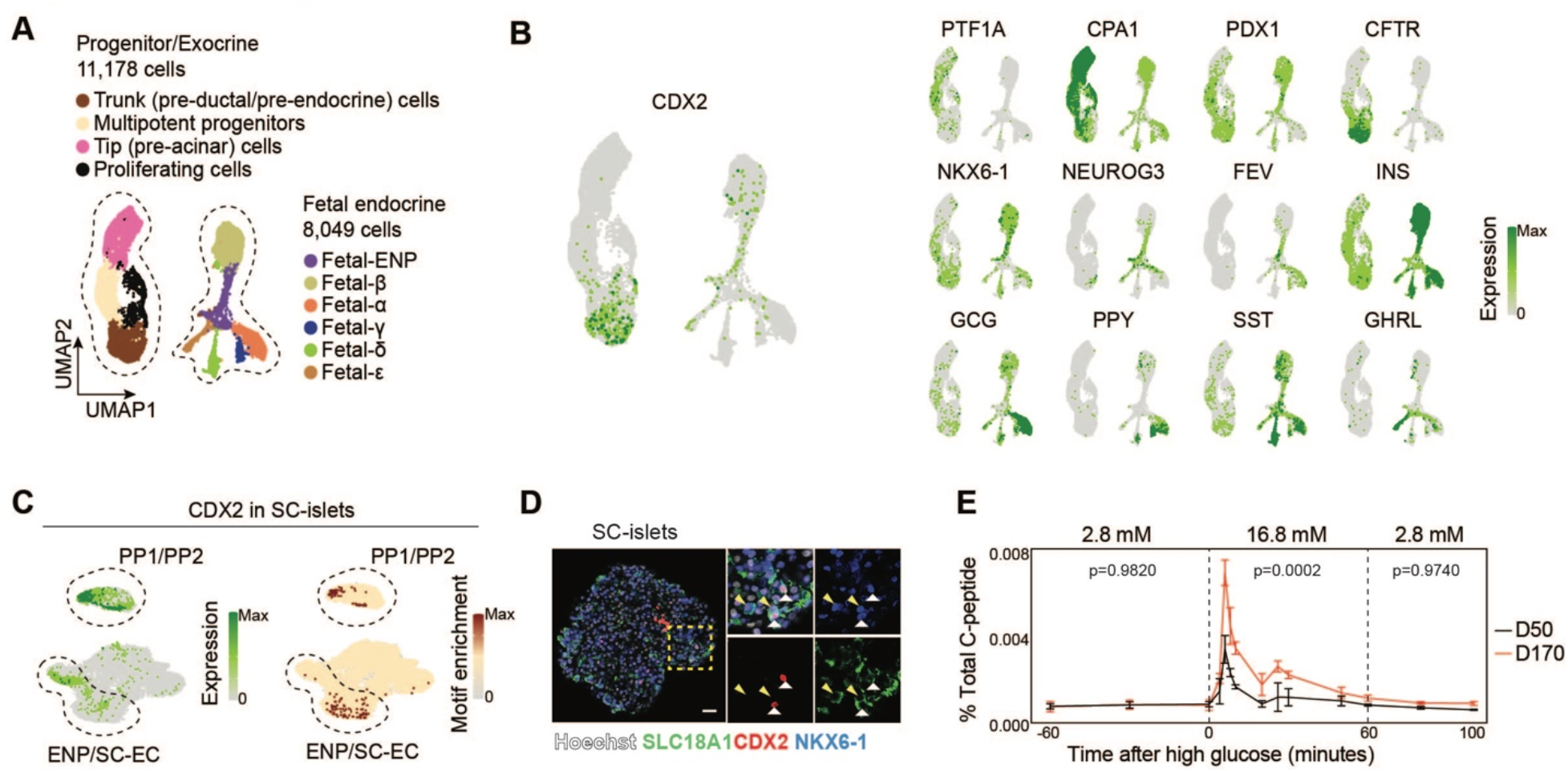
Identification of CDX2-expressing cells in human fetal pancreas. (A) UMAP embedding of transcriptome data from fetal human pancreas. Cluster identities were defined by expression of marker genes. (B) Expression of *CDX2* and cell type marker genes in fetal human pancreas. (C) *CDX2* expression (left) and CDX2 motif enrichment (right) in cell types during SC-islet differentiation. Dash circles highlight populations with high *CDX2* expression and transcriptional activity. (D) Representative immunofluorescent image for CDX2, NKX6-1, and SLC18A1 on SC-islets at D21. Arrowheads in insets indicate CDX2, NKX6-1 and SLC18A1 co-positive cells. Nuclei were labeled with Hoechst. Scale bar, 20 μm. (E) Human C-peptide secretion by SC-islets at D50 and D170 during perifusion with the indicated glucose concentrations (in mM). Data are shown as mean ± S.D. at the indicated time points. P-values were calculated using two-way ANOVA followed by Šidák-Holm’s multiple comparisons test for different days of differentiation in each time block.

**Supplemental Figure 5.**
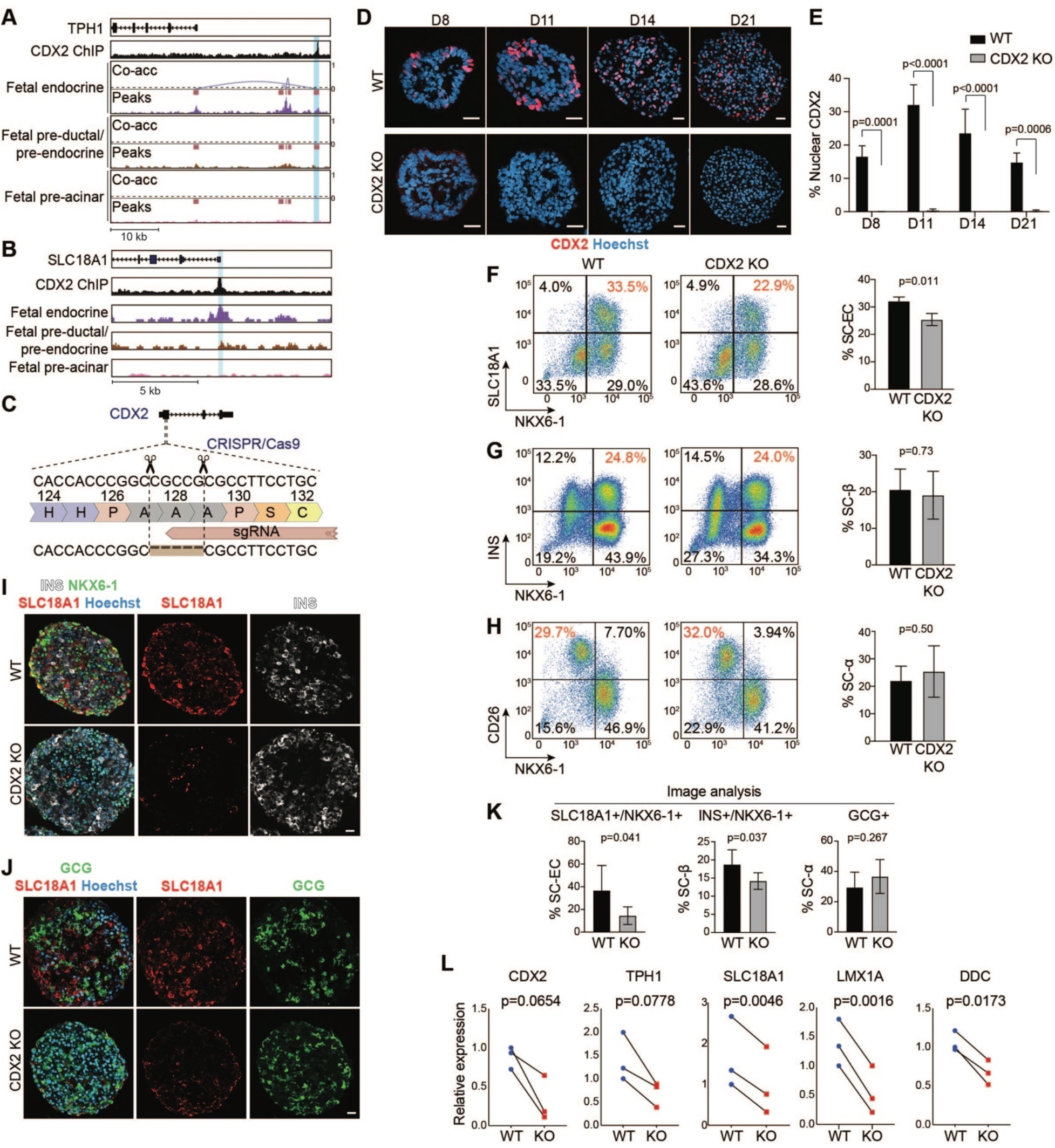
C*D*X2 inactivation in human pluripotent stem cells. (A, B) Genome browser tracks showing CDX2 ChIP-seq reads in SC-islets and aggregated ATAC reads in human fetal pancreatic endocrine, ductal and acinar cells at *TPH1* (A) and *SLC18A1* (B) gene loci. CDX2-bound cCREs are highlighted. All tracks are scaled to uniform 1×10^6^ read depth. Co-acc, co-accessibility scores between distal cCRE and transcriptional start site of *TPH1*. (C) Schematic showing generation of *CDX2* knockout (KO) H1 hPSCs. A 5-base-pair deletion was introduced into the first exon of *CDX2*, leading to a frameshift and premature translation termination. (D) Representative immunofluorescent images for CDX2 from wild type (WT) and *CDX2* KO cell aggregates at different days (D) of SC-islet differentiation. Nuclei were labeled with Hoechst. Scale bar, 20 μm. (E) Quantification of CDX2^+^ cells in WT and *CDX2* KO cell aggregates at different days of SC-islet differentiation. Data are shown as mean ± S.D. P-values were calculated using two-way ANOVA followed by Šidák-Holm’s multiple comparisons test for different days of differentiation. (F-H) Representative flow cytometry plots (left, percentage of population of interest in red) and quantifications (right) of SC-ECs (NKX6-1^+^/SLC18A1^+^, F), SC-β-cells (NKX6-1^+^/INS^+^, G), and SC-α-cells (NKX6-1^-^/CD26^+^, H) in WT and *CDX2* KO SC-islets at D21. Data are shown as mean ± S.D. (n = 3 independent differentiations). P-values were calculated by unpaired two-tailed t-test. (I) Representative immunofluorescent images for insulin (INS), NKX6-1 and SLC18A1 on WT and *CDX2* KO SC-islets at D21. Nuclei were labeled with Hoechst. Scale bar, 20 μm. (J) Representative immunofluorescent images for glucagon (GCG) and SLC18A1 on WT and *CDX2* KO SC-islets at D21. Nuclei were labeled with Hoechst. Scale bar, 20 μm. (K) Quantification of the percentage of SC-ECs (SLC18A1^+^/NKX6-1^+^), SC-β-cells (INS^+^/NKX6-1^+^) and SC-α-cells (GCG^+^) in SC-islets based on immunofluorescent staining in (I) and (J). Data are shown as mean ± S.D. (n = 3 independent differentiations). P-values were calculated by unpaired two-tailed t-test. (L) qPCR analysis of *CDX2* and serotonin synthesis genes in WT and *CDX2* KO SC-islets at D21. n = 100 SC-islets per group from 3 independent differentiations. P-values were calculated by two-tailed t-test.

**Supplemental Figure 6.**
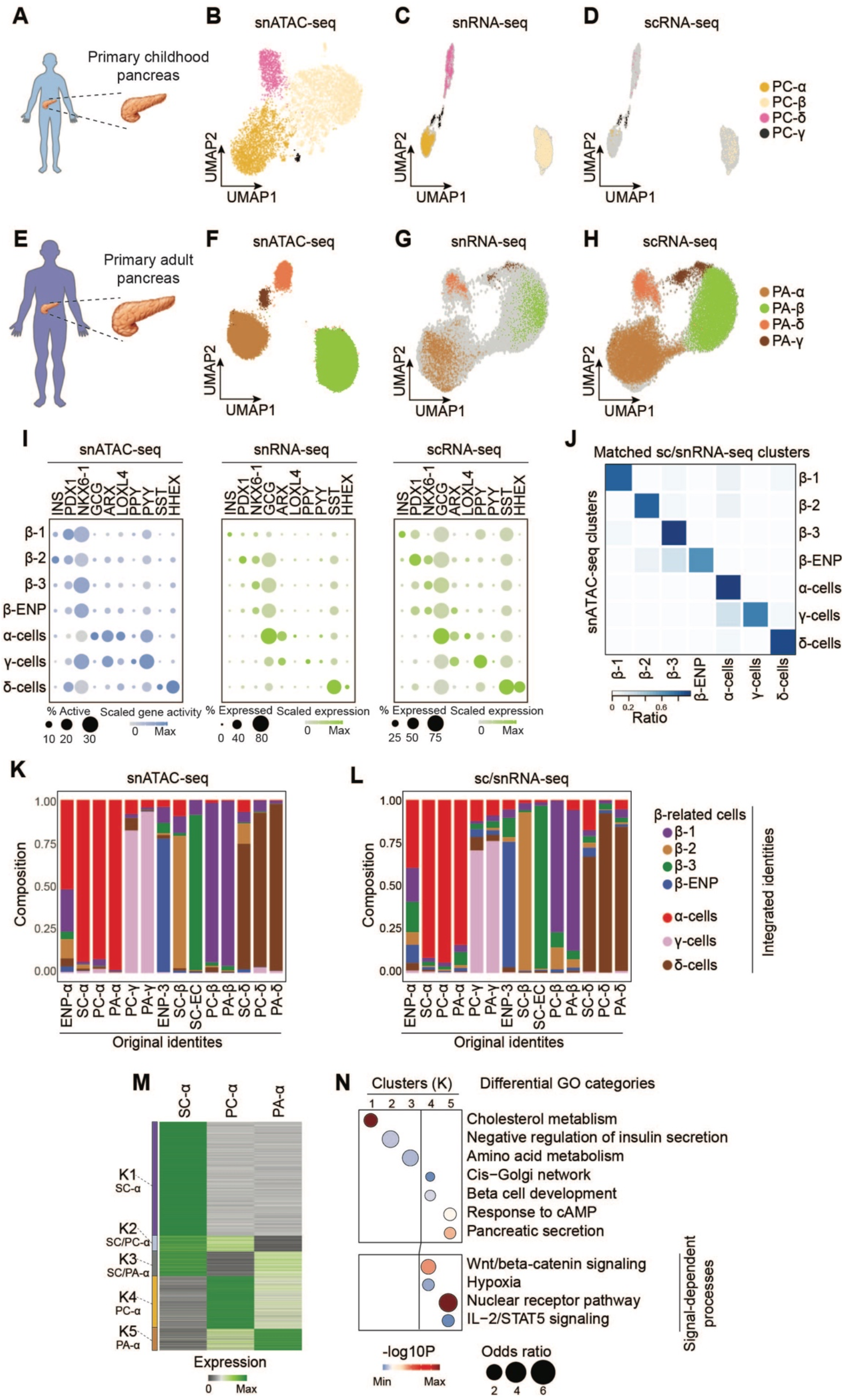
Integration of chromatin accessibility and transcriptome data from stem cell islets and primary pancreas. (A) Schematic showing source of cells used in single cell analyses of primary childhood human pancreas. (B-D) UMAP embedding of chromatin accessibility (B) and transcriptome (C and D) data from isolated nuclei (C) or whole cells (D) from childhood pancreas. Cluster identities were defined by promoter accessibility (snATAC-seq) or expression (sc/snRNA-seq) of marker genes. (E) Schematic showing source of cells used in single cell analyses of primary adult human pancreas. (F-H) UMAP embedding of chromatin accessibility (F) and transcriptome (G and H) data from isolated nuclei (G) or whole cells (H) from adult pancreas. Cluster identities were defined by promoter accessibility (snATAC-seq) or expression (sc/snRNA-seq) of marker genes. (I) Dot plots showing scaled average gene activity (left) or gene expression (middle and right). The color of each dot represents the average gene activity or expression level and the size of each dot the percentage of positive cells for each gene. (J) Heatmap showing ratio of cells with identities in sc/snRNA-seq (column) data matching identities in snATAC-seq (row) data. (K, L) Comparison of integrated cell type annotations based on snATAC-seq (K) and scRNA-seq (L) UMAP in Figure 6B,C to cell type annotations before data integration. (M) K-means clustering of genes with variable expression across α-related cell types (SC-α-cells, PC-α-cells, and PA-α-cells). Clusters were annotated and color-coded based on gene expression patterns. (N) Enriched gene ontology terms/pathways in each cluster. Significance (-log10 p-value) and odds ratio of the enrichments are represented by color and dot size, respectively.

**Supplemental Figure 7.**
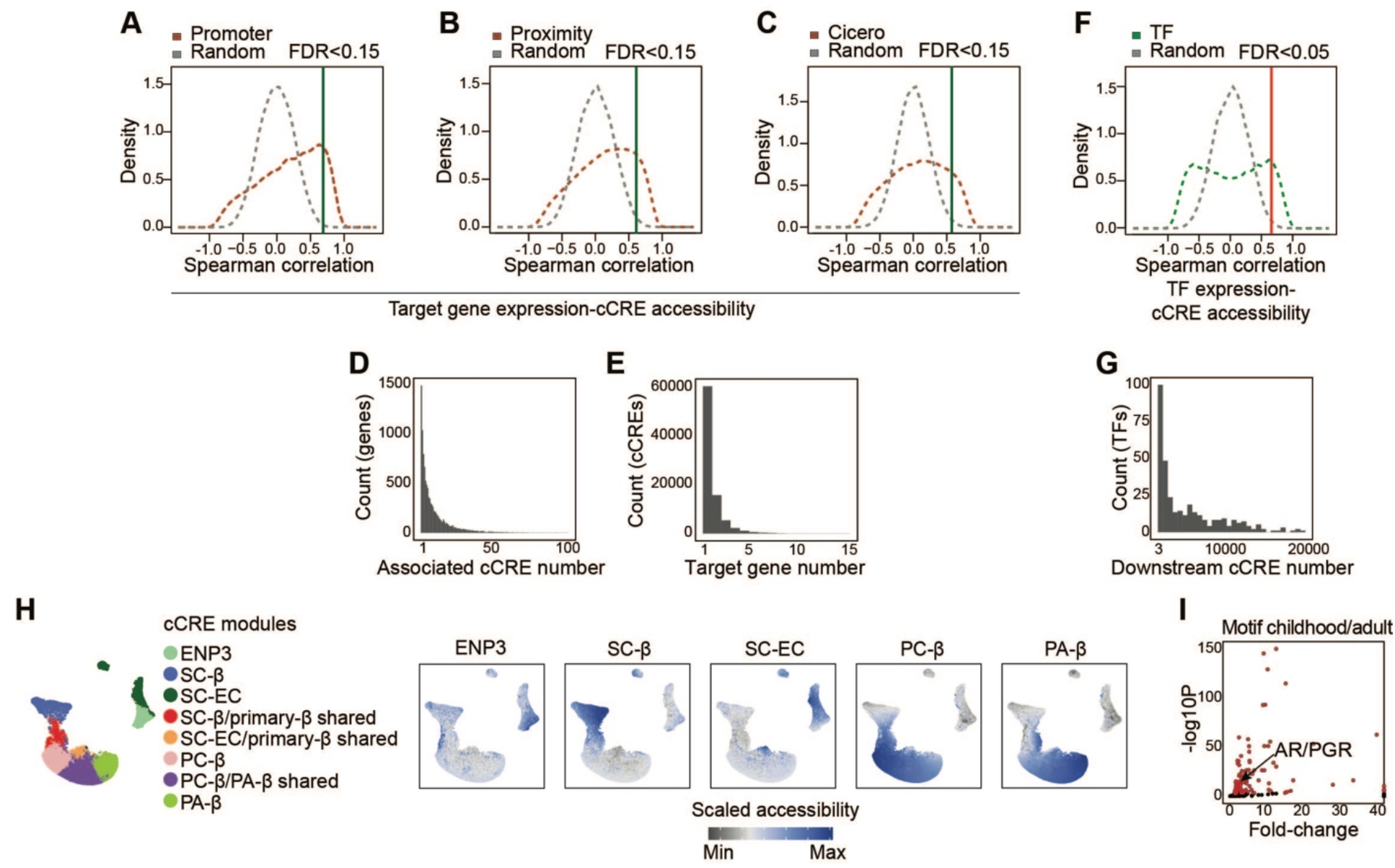
Building a gene regulatory network of β-cell maturation. (A-C) Identification of putative cCRE-gene pairs. cCREs were assigned to a gene if they are located (A) in promoter (± 500 bp) of target gene; (B) in proximity (± 50 kb) of the gene promoter; or (C) show co-accessibility with the gene promoter by Cicero analysis. A total of 167,618 positively correlated cCRE-gene pairs were identified using an empirically defined significance threshold of FDR<0.15. (D, E) Histogram showing characteristics of cCRE-gene pairs. Each target gene in the GRN is regulated by a mean of 13.6 cCREs (D) and each cCRE regulates a mean of 1.55 target genes (E). (F) Identification of putative TF-cCRE pairs. Putative TF-cCRE pairs were defined using motifmatchr. A total of 1,017,318 significantly correlated TF-cCRE pairs were identified using an empirically defined threshold of FDR<0.05. (G) Histogram showing characteristics of TF-cCRE pairs. Each TF has a mean of 2,698 predicted binding sites. (H) UMAP projections of scaled chromatin accessibility of all cCREs in each aggregated pseudo-cell. Pseudo-cells were aggregated based on cell type identities of primary cells and stem cell-derived populations before data integration. (I) Scatter plot showing fold-change and significance (-log10 p-value) of TF motif enrichment at chromatin sites with increased H3K27ac signals in childhood (<9 years) β-cells against a background of sites with increased H3K27ac signal in adult β-cells.

